# Conserved microRNA targeting reveals preexisting gene dosage sensitivities that shaped amniote sex chromosome evolution

**DOI:** 10.1101/118729

**Authors:** Sahin Naqvi, Daniel W. Bellott, Kathy S. Lin, David C. Page

## Abstract

Mammalian X and Y chromosomes evolved from an ordinary autosomal pair. Genetic decay of the Y led to X chromosome inactivation (XCI) in females, but some Y-linked genes were retained during the course of sex chromosome evolution, and many X-linked genes did not become subject to XCI. We reconstructed gene-by-gene dosage sensitivities on the ancestral autosomes through phylogenetic analysis of microRNA (miRNA) target sites and compared these preexisting characteristics to the current status of Y-linked and X-linked genes in mammals. Preexisting heterogeneities in dosage sensitivity, manifesting as differences in the extent of miRNA-mediated repression, predicted either the retention of a Y homolog or the acquisition of XCI following Y gene decay. Analogous heterogeneities among avian Z-linked genes predicted either the retention of a W homolog or gene-specific dosage compensation following W gene decay. Genome-wide analyses of human copy number variation indicate that these heterogeneities consisted of sensitivity to both increases and decreases in dosage. We propose a model of XY/ZW evolution incorporating such preexisting dosage sensitivities in determining the evolutionary fates of individual genes. Our findings thus provide a more complete view of the role of dosage sensitivity in shaping the mammalian and avian sex chromosomes, and reveal an important role for post-transcriptional regulatory sequences (miRNA target sites) in sex chromosome evolution.

## INTRODUCTION

The mammalian X and Y chromosomes evolved from a pair of ordinary autosomes over the past 300 million years (Lahn & Page, 1999). Only 3% of genes on the ancestral pair of autosomes survive on the human Y chromosome (Bellott et al., 2010; Skaletsky et al., 2003), compared to 98% on the X chromosome (Mueller et al., 2013). In females, one copy of the X chromosome is silenced by X inactivation (XCI); this silencing evolved on a gene-by-gene basis following Y gene loss and compensatory X upregulation (Berletch et al., 2015; Jegalian & Page, 1998; Ross et al., 2005; Tukiainen et al., 2017), and some genes escape XCI in humans (Carrel & Willard, 2005) and other mammals (Yang, Babak, Shendure, & Disteche, 2010). In mammals, dosage compensation, which refers to any mechanism restoring ancestral dosage following gene loss from the sex-specific chromosome, thus consists of both X upregulation following Y gene loss and the subsequent acquisition of XCI.

In parallel, the avian Z and W sex chromosomes evolved from a different pair of autosomes than the mammalian X and Y chromosomes (Bellott et al., 2010; Nanda et al., 1999; Ross et al., 2005). Decay of the female-specific W chromosome was similarly extensive, but birds did not evolve a large-scale inactivation of Z-linked genes analogous to XCI in mammals (Itoh et al., 2007). Dosage compensation, as measured by a male/female expression ratio close to 1, has been observed for some Z-linked genes in some tissues. (Mank & Ellegren, 2009; Uebbing et al., 2015; Zimmer, Harrison, Dessimoz, & Mank, 2016). Thus, genes previously found on the ancestral autosomes that gave rise to the mammalian or avian sex chromosomes have undergone significant changes in gene dosage. In modern mammals, these molecular events have resulted in three classes of ancestral X-linked genes representing distinct evolutionary fates: those with a surviving Y homolog, those with no Y homolog and subject to XCI, and those with no Y homolog but escaping XCI. In birds, two clear classes of ancestral Z-linked genes have arisen: those with or without a W homolog, with additional heterogeneity among Z-linked genes without a W homolog as a result of gene-specific dosage compensation. Identifying gene-by-gene properties that distinguish classes of X- and Z-linked genes is thus crucial to understanding the selective pressures underlying the molecular events of mammalian and avian sex chromosome evolution.

Emerging evidence suggests a role for gene dosage sensitivity in mammalian and avian sex chromosome evolution. X- and Z-linked genes with surviving homologs on the mammalian Y or avian W chromosomes are enriched for important regulatory functions and predictors of haploinsufficiency compared to those lacking Y or W homologs (Bellott et al., 2014, 2017); similar observations have been made in fish (White, Kitano, & Peichel, 2015) and Drosophila (Kaiser, Zhou, & Bachtrog, 2011). Human X- and chicken Z-linked genes that show the strongest signatures of dosage compensation in either lineage also show signs of dosage sensitivity as measured by membership in large protein complexes (Pessia, Makino, Bailly-Bechet, McLysaght, & Marais, 2012) or evolutionary patterns of gene duplication and retention (Zimmer et al., 2016). Despite these advances, little is known regarding selective pressures resulting from sensitivity to dosage increases, as these studies either focused on haploinsufficiency or employed less direct predictors of dosage sensitivity. Furthermore, it is not known whether heterogeneities in dosage sensitivity among classes of sex-linked genes were acquired during sex chromosome evolution, or predated the emergence of sex chromosomes, as there has been no explicit, systematic reconstruction of dosage sensitivity on the ancestral autosomes that gave rise to the mammalian and avian sex chromosomes.

To assess the role of preexisting dosage sensitivities in XY and ZW evolution, we sought to employ a measure of dosage sensitivity that could be 1) demonstrably informative with respect to sensitivity to dosage increases, and 2) explicitly reconstructed on the ancestral autosomes. We focused on regulation by microRNAs (miRNAs), small noncoding RNAs that function as tuners of gene dosage by lowering target mRNA levels through pairing to the 3 ̀ untranslated region (UTR) (Bartel, 2009). The repressive nature of miRNA targeting is informative with respect to sensitivity to dosage increases, allowing for a more complete understanding of the role of dosage sensitivity in sex chromosome evolution. Both miRNAs themselves and their complementary target sites can be preserved over millions of years of vertebrate evolution, facilitating the reconstruction of miRNA targeting on the ancestral autosomes through cross-species sequence alignments. As miRNA targeting occurs post-transcriptionally, reconstruction of its ancestral state is decoupled from transcriptional regulatory mechanisms such as XCI that evolved following X-Y differentiation.

## RESULTS

### Analysis of human copy number variation indicates conserved microRNA targeting of genes sensitive to dosage increases

We first sought to determine whether conserved targeting by microRNAs (miRNAs) correlates with sensitivity to dosage increases across the human genome. To estimate pressure to maintain miRNA targeting, we used published probabilities of conserved targeting (P_CT_ scores) for each gene-miRNA interaction in the human genome. The P_CT_ score reflects an estimate of the probability that a given gene-miRNA interaction is conserved due to miRNA targeting, obtained by calculating the conservation of the relevant miRNA target sites relative to the conservation of the entire 3 ̀ UTR (Friedman, Farh, Burge, & Bartel, 2009). In this manner, the P_CT_ score intrinsically controls for differences in background conservation and sequence composition, both of which vary widely among 3 ̀UTRs due to differing rates of expression divergence and/or sequence evolution. We refer to these P_CT_ scores as “miRNA conservation scores” in the remainder of the text.

A recent study reported a correlation between these miRNA conservation scores and predicted haploinsufficiency (Pinzón et al., 2016), indicating that conserved miRNA targeting broadly corresponds to dosage sensitivity. However, such a correlation does not isolate the effects of sensitivity to dosage increases, which we expect to be particularly important in the context of miRNA targeting. We reasoned that genes for which increases in dosage are deleterious should be depleted from the set of observed gene duplications in healthy human individuals. We used a catalogue of rare genic copy number variation among 59,898 control human exomes (Exome Aggregation Consortium, ExAC)(Ruderfer et al., 2016) to classify autosomal protein-coding genes as exhibiting or lacking duplication or deletion in healthy individuals (see Methods). We compared duplicated and non-duplicated genes with the same deletion status in order to control for differences in sensitivity to underexpression. We found that non-duplicated genes have significantly higher miRNA conservation scores than duplicated genes, irrespective of deletion status (Figure 1A,B). Non-deleted genes also have significantly higher scores than deleted genes irrespective of duplication status (Supplemental Figure S1), but duplication status has a greater effect on miRNA conservation scores than does deletion status (blue vs. orange boxes, Figure 1C). Thus, conserved miRNA targeting is a feature of genes sensitive to changes in gene dosage in humans and is especially informative with regards to sensitivity to dosage increases, consistent with the known role of miRNAs in tuning gene dosage by lowering target mRNA levels.

**Figure 1:**
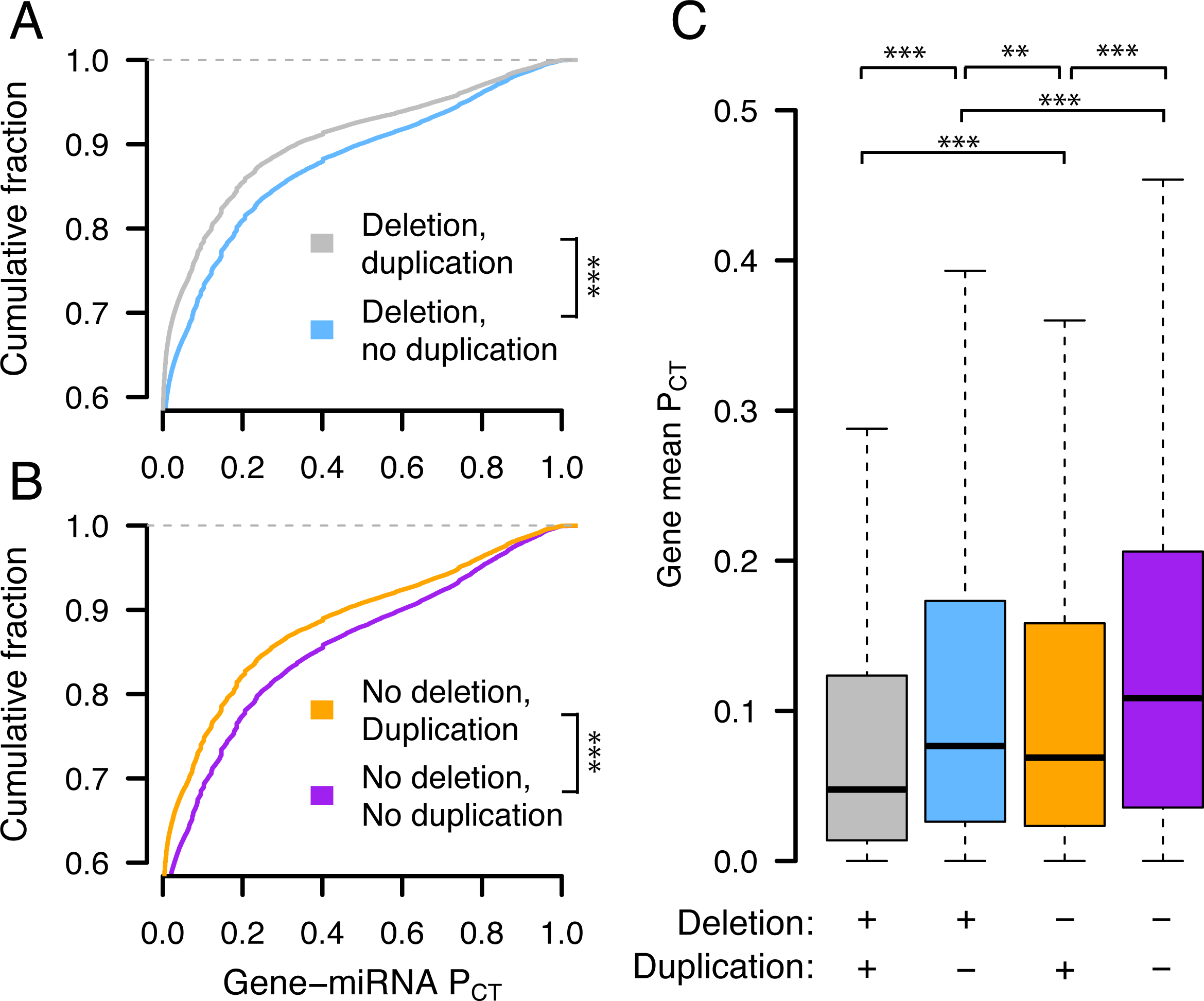
Conserved miRNA targeting of autosomal genes stratified by copy number variation in 59,898 human exomes. Probabilities of conserved targeting (P_CT_) of all gene-miRNA interactions involving non-duplicated and duplicated genes, further stratified as (A) deleted (grey, n = 69,339 interactions from 4,118 genes; blue, n = 80,290 interactions from 3,976 genes) or (B) not deleted (orange, n = 51,514 interactions from 2,916 genes; purple, n = 72,826 interactions from 3,510 genes). *** p < 0.001, two-sided Kolmogorov-Smirnov test. (C) Mean gene-level P_CT_ scores. ** p < 0.01, *** p < 0.001, two-sided Wilcoxon rank-sum test.

### X-Y pairs and X-inactivated genes have higher miRNA conservation scores than X escape genes

We next assessed whether the three classes of X-linked genes differ with respect to dosage sensitivity as inferred by conserved miRNA targeting. To delineate these classes, we began with the set of ancestral genes reconstructed through cross-species comparisons of the human X chromosome and orthologous chicken autosomes (Bellott et al., 2014, 2017, 2010; Hughes et al., 2012; Mueller et al., 2013). We designated ancestral X-linked genes with a surviving human Y homolog (Skaletsky et al., 2003) as X-Y pairs and also considered the set of X-linked genes with a surviving Y homolog in any of eight mammals (Bellott et al., 2014) to increase the phylogenetic breadth of findings regarding X-Y pairs. A number of studies have catalogued the inactivation status of X-linked genes in various human tissues and cell-types. We used a meta-analysis that combined results from three studies by assigning a “consensus” X-inactivation status to each gene (Balaton, Cotton, & Brown, 2015) to designate the remainder of ancestral genes lacking a Y homolog as subject to or escaping XCI. In summary, we classified genes as either: 1) X-Y pairs, 2) lacking a Y homolog and subject to XCI (X-inactivated), or 3) lacking a Y homolog but escaping XCI (X escape).

We found that human X-Y pairs have the highest miRNA conservation scores, followed by X-inactivated and finally X escape genes (Figure 2A,B). The expanded set of X-Y pairs across eight mammals also has significantly higher miRNA conservation scores than ancestral X-linked genes with no Y homolog (Supplemental Figure S2). Observed differences between miRNA conservation scores are not driven by distinct subsets of genes in each class, as indicated by gene resampling with replacement (Supplemental Figure S3). The decrease in miRNA conservation scores of X escape genes relative to X-inactivated genes and X-Y pairs is not driven by genes that escape XCI variably across individuals (Supplemental Figure S4), and was consistent even when including ambiguous genes as either X-inactivated or X escape genes (Supplemental Figure S5). We also verified that these differences were not driven by artificially inflated or deflated conservation scores of certain target sites due to non-uniformity in 3 ̀ UTR conservation (Methods, Supplemental Figure S6).

**Figure 2.**
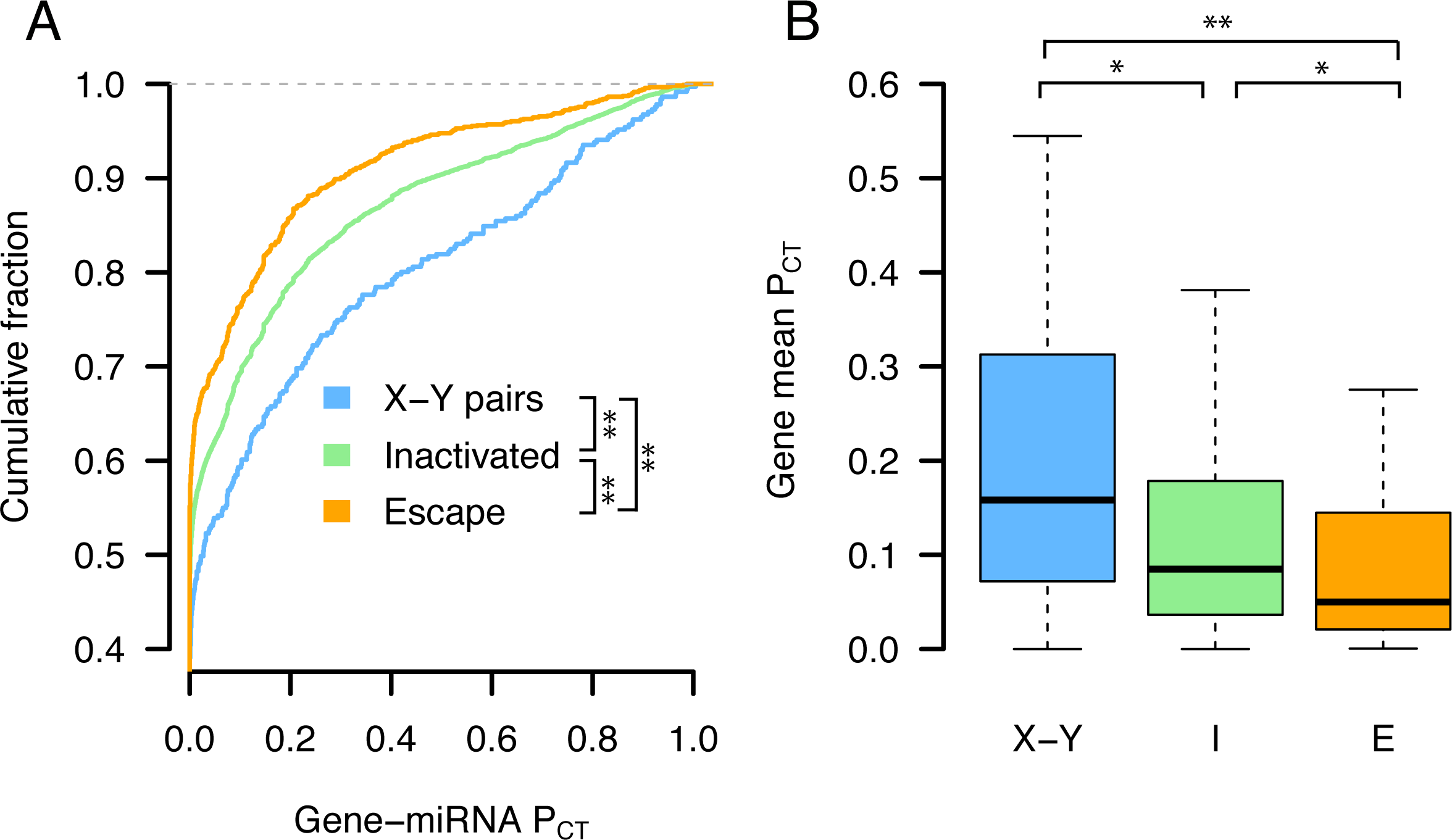
X-Y pairs and X-inactivated genes have higher miRNA conservation scores than X escape genes. P_CT_ score distributions of all gene-miRNA interactions involving (A) human X-Y pairs (n = 371 interactions from 15 genes), X-inactivated genes (n = 6,743 interactions from 329 genes), and X escape genes (n = 1,037 interactions from 56 genes). ** p < 0.01, two-sided Kolmogorov-Smirnov test. (B) Mean gene-level P_CT_ scores. * p < 0.05, ** p < 0.01, two-sided Wilcoxon rank-sum test.

Finally, we assessed whether miRNA conservation scores distinguish the three classes by providing additional information not accounted for by known factors (Bellott et al., 2014) influencing evolutionary outcomes. We used logistic regression to model, for each gene, the probability of falling into each of the three classes (X-Y pair, X-inactivated, or X escape) as a linear combination of haploinsufficiency probability (pHI) (Huang, Lee, Marcotte, & Hurles, 2010), human expression breadth (GTEx Consortium, 2015), purifying selection, measured by the ratio of non-synonymous to synonymous substitution rates (dN/dS) between human and mouse orthologs (Yates et al., 2016), and mean gene-level miRNA conservation scores. We note that pHI is a score composed of several genic features, one of which is the number of protein-protein interactions, consistent with the idea that members of large protein complexes tend to be dosage-sensitive (Papp, Pal, & Hurst, 2003; Pessia et al., 2012). Removing either miRNA conservation or pHI as predictors from the full model resulted in inferior model fits as measured by Aikake’s information criterion (AIC) (full model, AIC 321.5; full model minus miRNA conservation, AIC 327.7; full model minus pHI, AIC 327.3; higher AIC indicates inferior model). Therefore, miRNA conservation and pHI contribute independent information that distinguishes the 3 classes of X-linked genes. Based on our analyses of autosomal copy number variation (Figure 1), we attribute this independence to the fact that miRNA conservation scores are most informative with regards to sensitivity to dosage increases. Taken together, these results indicate significant heterogeneity in dosage sensitivity, as inferred by miRNA target site conservation, among the three classes of ancestral X-linked genes: X-Y pairs are the most dosage-sensitive, while X-inactivated genes are of intermediate dosage sensitivity, and X escape genes are the least dosage-sensitive.

### Heterogeneities in X-linked miRNA targeting were present on the ancestral autosomes

We next asked whether differences in miRNA targeting were present on the ancestral autosomes that gave rise to the mammalian X and Y chromosomes. To reconstruct the ancestral state of miRNA targeting, we first focused on miRNA target sites in the 3 ̀ UTR of human orthologs that align with perfect identity to a site in the corresponding chicken ortholog; these sites were likely present in the common ancestor of mammals and birds (Figure 3A,B). We found that X-Y pairs have the most human-chicken conserved target sites, followed by X-inactivated genes, and then X escape genes (Figure 3C, top). Unlike the miRNA conservation scores used earlier, this metric does not account for background conservation; we therefore estimated the background conservation of each 3 ̀ UTR using shuffled miRNA family seed sequences (see Methods). X-Y pairs, X-inactivated genes, and X escape genes do differ significantly with respect to background conservation (data not shown), but these differences cannot account for the observed differences in true human-chicken conserved sites (Figure 3C, bottom). We observed similar results for the expanded set of X-Y pairs across 8 mammals (Supplemental Figure S7A).

**Figure 3.**
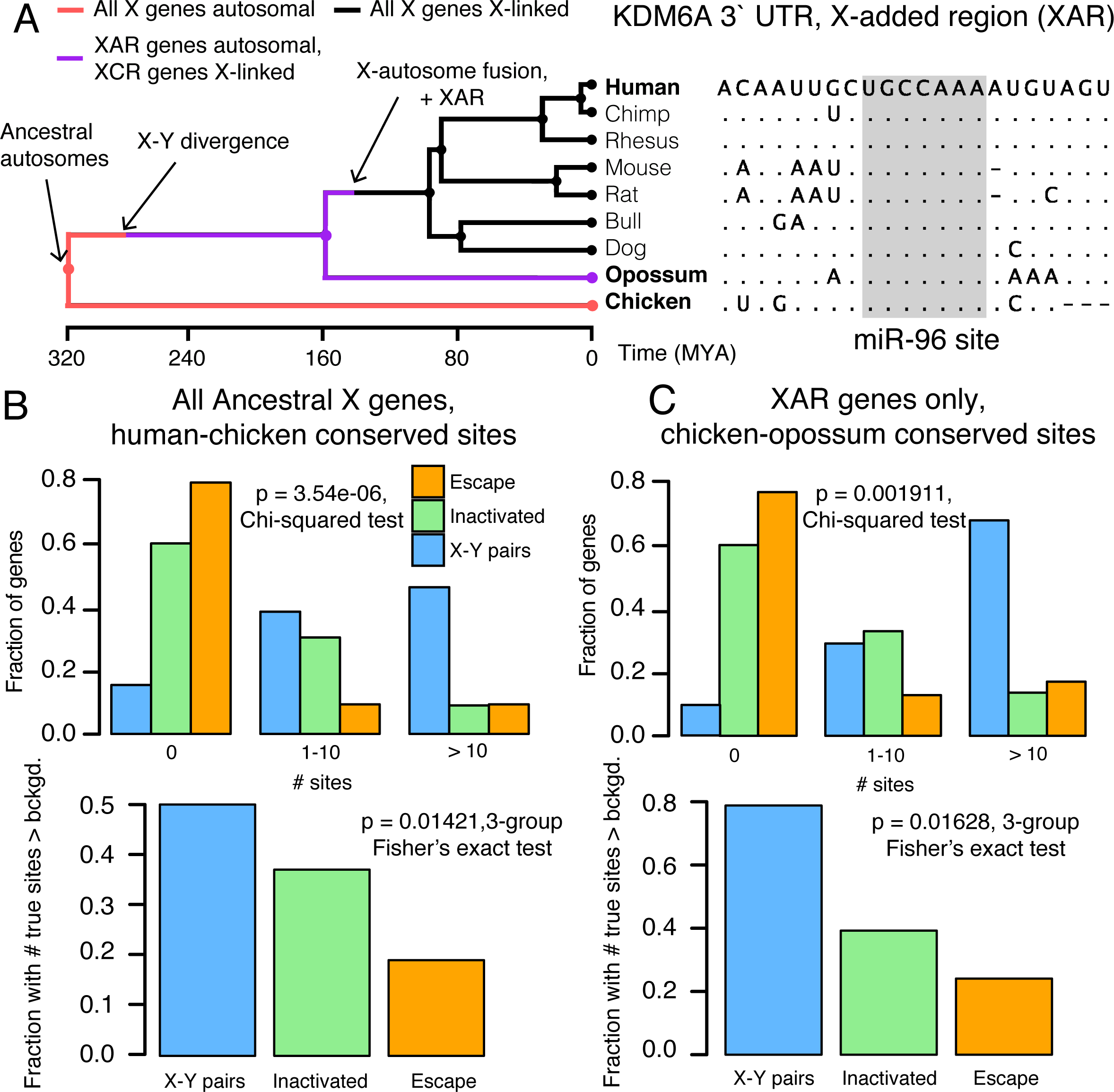
Heterogeneities in X-linked miRNA targeting were present on the ancestral autosomes. (A) Example reconstruction of an ancestral miR-96 target site in the 3 ̀ UTR of KDM6A, an X-linked gene in the X-added region (XAR) with a surviving Y homolog. Dots in non-human species indicate identity with the human sequence, dashes gaps indicate gaps in the multiple sequence alignment. (B) Distributions of sites conserved between 3 ̀ UTRs of human and chicken orthologs (top) or comparisons to background expectation (bottom, see Methods) for human X-Y pairs (n = 16), X-inactivated genes (n = 251), and X escape genes (n = 42). (C)Statistics as in (B), but using sites conserved between chicken and opossum 3 ̀ UTRs only for genes in the XAR; X-Y pairs (n = 11), X-inactivated genes (n = 58), and X escape genes (n = 27).

Differences in the number of human-chicken conserved sites among the three classes of X-linked genes could be explained by heterogeneity in miRNA targeting present on the ancestral autosomes, or by ancestral homogeneity followed by different rates of target site loss during or following X-Y differentiation. To distinguish between these two possibilities, we took advantage of previous reconstructions of human sex chromosome evolution (Figure 3A) (Bellott et al.,2014), which confirmed that, following the divergence of placental mammals from marsupials, an X-autosome chromosomal fusion generated the X-added region (XAR) (Watson, Spencer, Riggs, & Graves, 1990). Genes on the XAR are therefore X-linked in placental mammals, but autosomal in marsupials such as the opossum. We limited our analysis to genes in the XAR and target sites conserved between orthologous chicken and opossum 3 ̀ UTRs, ignoring site conservation in humans; these sites were likely present in the common ancestor of mammals and birds, and an absence of such sites cannot be explained by site loss following X-Y differentiation. We observed the same pattern as with the human-chicken conserved sites, both before and after accounting for background 3 ̀ UTR conservation (Figure 3D, three gene classes; Supplemental Figure S7B, X-Y pairs across 8 mammals). These results demonstrate that the autosomal precursors of X-Y pairs and X-inactivated genes were subject to more miRNA-mediated regulation than X escape genes. Combined with our earlier results, we conclude that present-day heterogeneities in dosage sensitivity on the mammalian X chromosome existed on the ancestral autosomes from which it derived.

### Z-W pairs have higher miRNA conservation scores than other ancestral Z-linked genes

We next assessed whether classes of avian Z-linked genes, those with and without a W homolog, show analogous heterogeneities in sensitivity to dosage increases. We used the set of ancestral genes reconstructed through cross-species comparisons of the avian Z chromosome and orthologous human autosomes and focused on the set of Z-W pairs identified by sequencing of the chicken W chromosome (Bellott et al., 2017, 2010). To increase the phylogenetic breadth of our comparisons, we also included candidate Z-W pairs obtained through comparisons of male and female genome assemblies (4 species set) or inferred by read-depth changes in female genome assemblies (14 species set, see Methods for details) (Zhou et al., 2014). The more complete 3 ̀ UTR annotations in the human genome relative to chicken allow for a more accurate assessment of conserved miRNA targeting. Accordingly, we analyzed the 3 ̀ UTRs of the human orthologs of chicken Z-linked genes.

We found that the human orthologs of Z-W pairs have higher miRNA conservation scores than the human orthologs of other ancestral Z genes (Figure 4A, B). Differences in miRNA conservation scores between Z-W pairs and other ancestral Z genes remained significant when considering the expanded sets of Z-W pairs across four and 14 avian species (Supplemental Figure S8). These differences are not driven by distinct subsets of genes, as indicated by gene resampling with replacement (Supplemental Figure S9), and cannot be accounted for by within-UTR variation in regional conservation (Supplemental Figure S10). Logistic regression models indicate that miRNA conservation scores provide additional information not captured by known factors (Bellott et al., 2017) influencing survival of W-linked genes (full model model, AIC 127.1; full model minus miRNA conservation, AIC 137.8; full model minus pHI 132.7; higher AIC indicates inferior model). Together, these results indicate that Z-linked genes with a surviving W homolog are more sensitive to changes in dosage -- both increases and decreases -- than are genes without a surviving W homolog.

**Figure 4.**
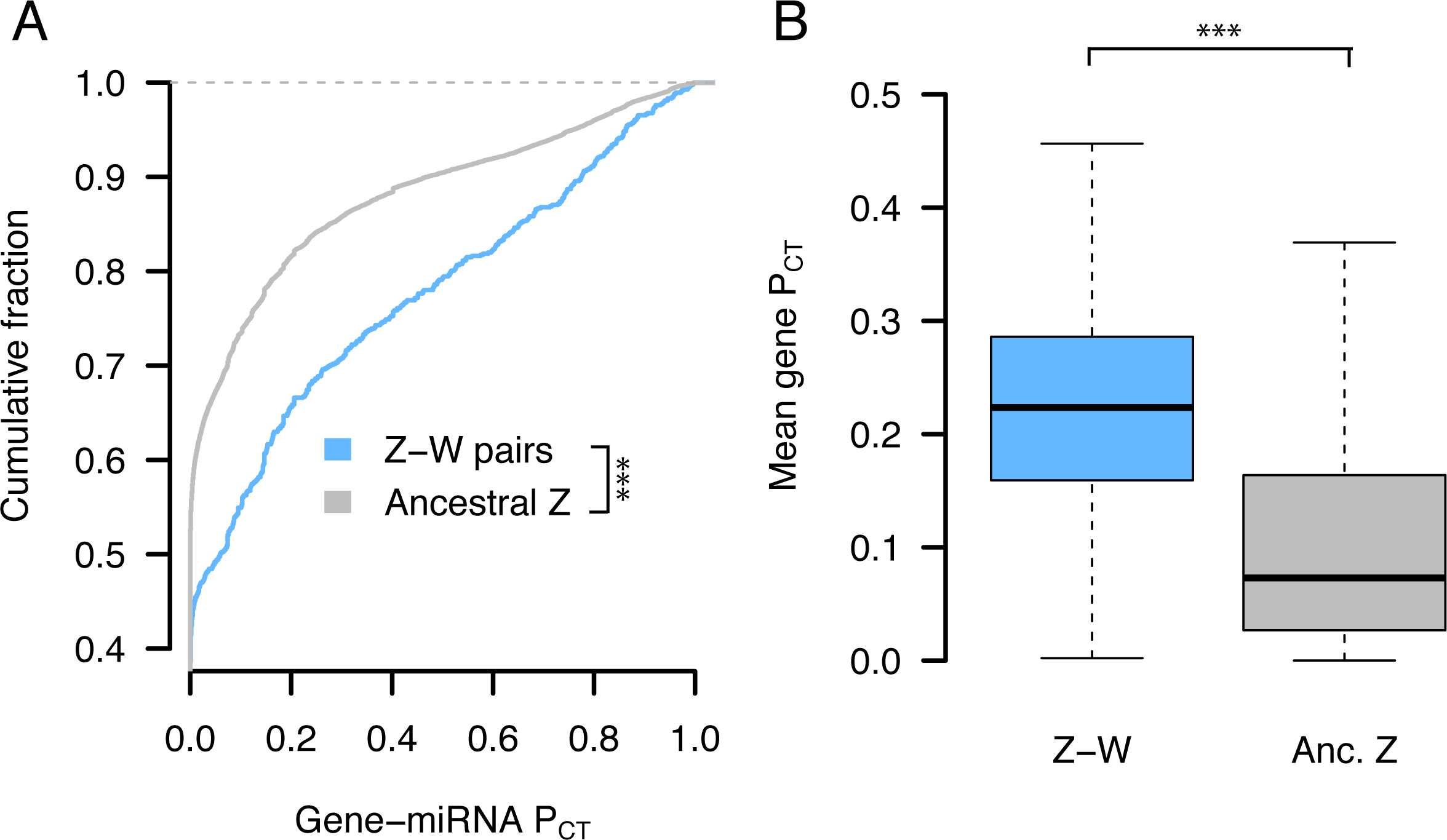
Z-W pairs have higher miRNA conservation scores than other ancestral Z-linked genes. P_CT_ score distributions of all gene-miRNA interactions involving the human orthologs of (A) chicken Z-W pairs (n = 832 interactions from 28 genes) and other ancestral Z genes (n = 16,692 interactions from 657 genes). *** p < 0.001, two-sided Kolmogorov-Smirnov test. (B) Mean gene-level P_CT_ scores. *** p < 0.001, two-sided Wilcoxon rank-sum test.

While there are two clear classes of Z-linked genes -- those with or without a W homolog -- studies of Z-linked gene expression have suggested additional heterogeneity among Z-linked genes without a W homolog due to gene-specific dosage compensation (Mank & Ellegren, 2009; Uebbing et al., 2015; Zimmer et al., 2016). If Z-linked genes with no W homolog exist upon a continuum from non-compensated to dosage-compensated, those that are more compensated should have more conserved miRNA target sites, reflective of greater dosage sensitivity. We quantified the dosage compensation by using RNA sequencing data (Marin et al., 2017) to compare, in 4 somatic tissues, the chicken male/female expression ratio to the analogous ratio in human and anolis (see Methods). In the brain, kidney, and liver, Z-linked genes with no W homolog and higher mean miRNA conservation scores had male/female expression ratios closer to 1 (Supplemental Figure S11). Thus, in addition to the above-described differences between Z-linked genes with or without a W homolog, Z-linked genes with no W homolog but with more effective dosage compensation have more conserved miRNA target sites than non-compensated genes.

### Heterogeneities in Z-linked miRNA targeting were present on the ancestral autosomes

We next asked whether differences in miRNA targeting between Z-W pairs and other ancestral Z-linked genes were present on the ancestral autosomes that gave rise to the avian Z and W chromosomes. We found that chicken Z-W pairs have more human-chicken-conserved miRNA target sites than their Z-linked counterparts without surviving W homologs, both before (Figure 5C, top) and after (Figure 5C, bottom) accounting for the background conservation of each individual 3 ̀ UTR. To confirm that these differences represent ancestral heterogeneity rather than differential site loss during or following Z-W differentiation, we instead considered the number of sites conserved between human and anolis lizard, which diverged from birds prior to Z-W differentiation (Figure 5A). Chicken Z-W pairs contain an excess of human-anolis conserved miRNA target sites, both before (Figure 5D, top) and after (Figure 5D, bottom) accounting for the background conservation of each individual 3 ̀ UTR. We observed similar results with the predicted four-species (Supplemental Figure S12) and 14-species (Supplemental Figure S13) sets of Z-W pairs. Thus, the autosomal precursors of avian Z-W pairs were subject to more miRNA-mediated regulation than the autosomal precursors of Z-linked genes that lack a W homolog. Furthermore, in the liver and brain, Z-linked genes with no W homolog with an excess of human-chicken-conserved miRNA sites had male/female expression ratios closer to 1, implying more effective dosage compensation (Supplementary Figure S14). Together, these results indicate heterogeneity in dosage sensitivity among genes on the ancestral autosomes that gave rise to the avian Z chromosome.

**Figure 5.**
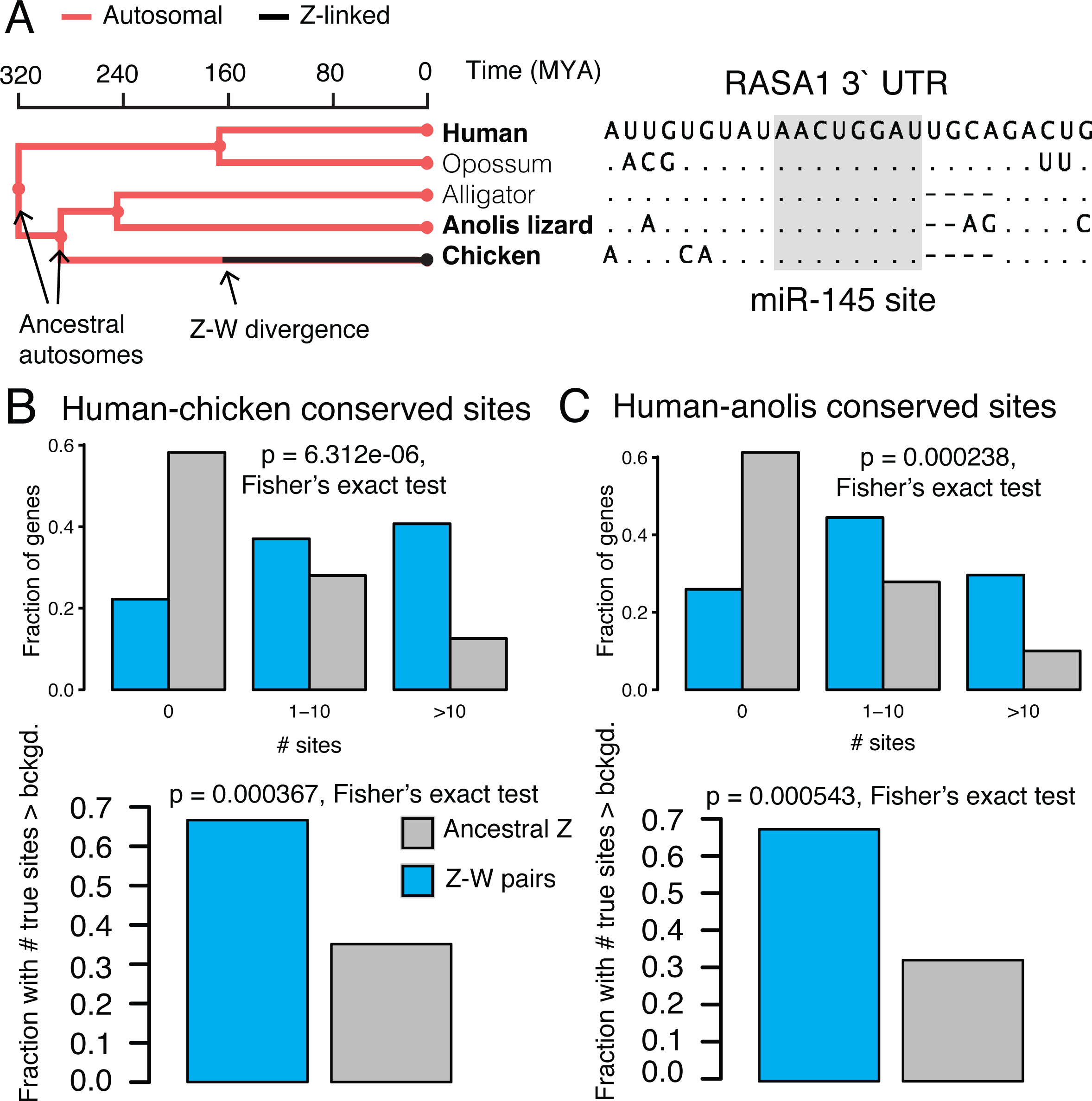
Heterogeneities in Z-linked miRNA targeting were present on the ancestral autosomes. (A) Example reconstruction of an ancestral miR-145 target site in the 3 ̀ UTR of RASA1, a Z-linked gene with a surviving W homolog. Example of 3 ̀ UTR sequence alignment for RASA1, a Z-linked gene with a surviving W homolog, with a target site for miR-145 highlighted in gray. (B) Numbers of sites conserved between 3 ̀ UTRs of human and chicken orthologs (top) or comparisons to background expectation (bottom) for chicken Z-W pairs (n = 27) and other ancestral Z genes (n = 578). (C) Statistics as in (B), but using sites conserved between human and anolis 3 ̀ UTRs.

### Analyses of experimental datasets validate miRNA target site function

Our results to this point, which indicate preexisting heterogeneities in dosage constraints among X- or Z-linked genes as inferred by predicted miRNA target sites, lead to predictions regarding the function of these sites in vivo. To test these predictions, we turned to publically available experimental datasets consisting both of gene expression profiling following transfection or knockout of individual miRNAs, and of high-throughput crosslinking-immunoprecipitation (CLIP) to identify sites that bind Argonaute in vivo (see Methods). If the above-studied sites are effective in mediating target repression, targets of an individual miRNA should show increased expression levels or Argonaute binding following miRNA transfection, and decreased expression levels following miRNA knockout. Together, our analyses of publically available datasets fulfilled these predictions, validating the function of these sites in multiple cellular contexts and species (Figure 6). From the gene expression profiling data, we observed results consistent with effective targeting by a) eleven different miRNA families in human HeLa cells (Supplemental Figure S15), b) four different miRNAs in human HCT116 and HEK293 cells (Supplemental Figure S16), and c) miR-155 in mouse B and Th1 cells (Supplemental Figure S17). In the CLIP data, the human orthologs of X- or Z-linked targets of miR-124 are enriched for Argonaute-bound clusters that appear following miR-124 transfection, while a similar but non-significant enrichment is observed for miR-7 (Supplemental Figure S18). Thus, conserved miRNA target sites used to infer dosage constraints on X-linked genes and the autosomal orthologs of Z-linked genes can effectively mediate target repression in living cells.

**Figure 6.**
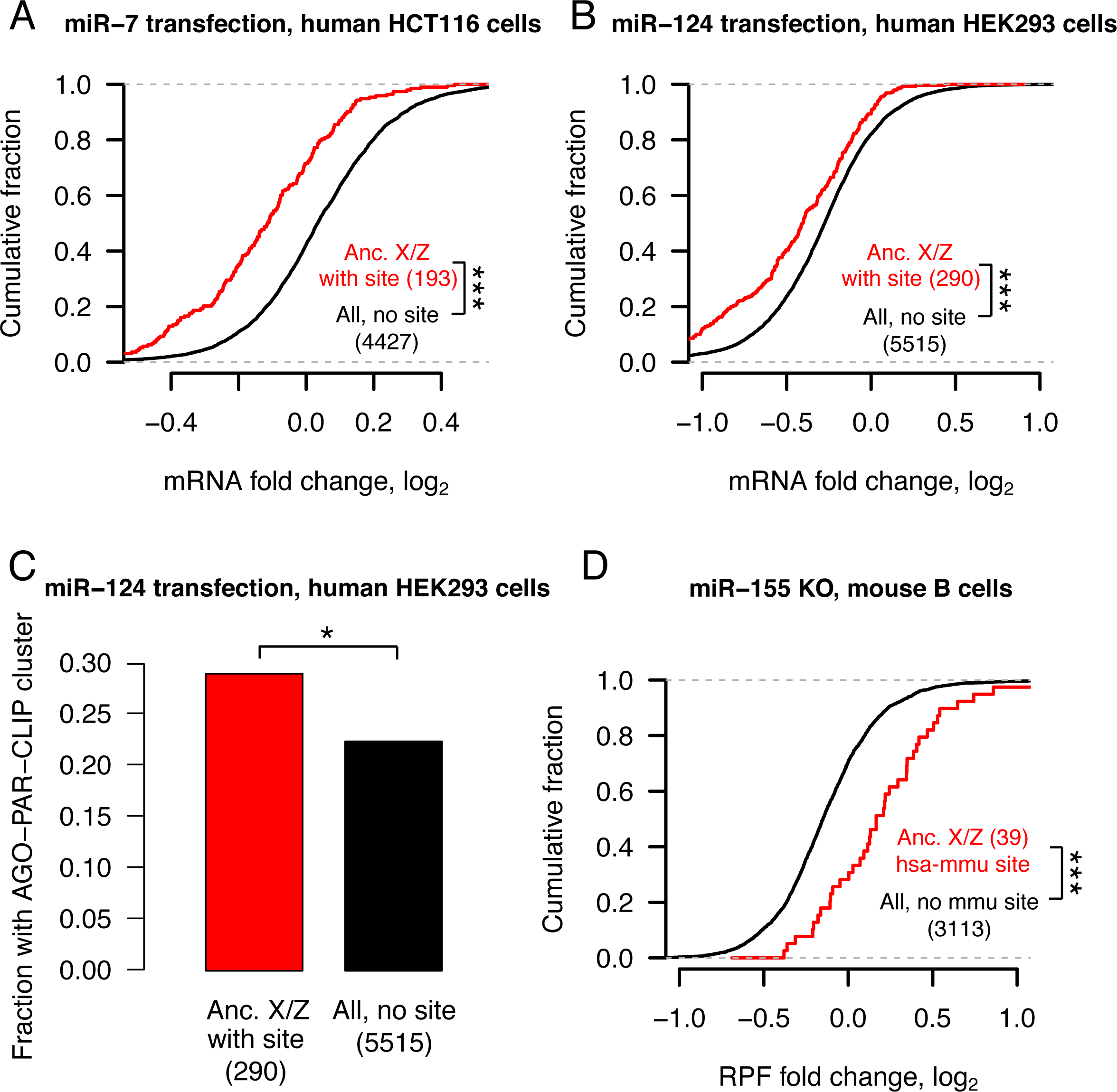
Analyses of experimental datasets validate miRNA target site function. Responses to transfection (A,B,C) or knockout (D) of indicated miRNAs in human (A,B,C) or mouse (D) cell-types. Each panel depicts corresponding changes in mRNA levels (A,B), in fraction of Argonaute-bound genes (C), and in mRNA stability and translational efficiency as measured by ribosome protected fragments (RPF, D). In each case, X-linked genes and the human orthologs of Z-linked genes containing target sites with an assigned P_CT_ score (red) for the indicated miRNA were compared to all expressed genes lacking target sites (black); gene numbers are indicated in parentheses. (A,B,D) *** p < 0.001, two-sided Kolmogorov-Smirnov test. (C) * p < 0.05, two-sided Fisher’s exact test.

## DISCUSSION

Here, through the evolutionary reconstruction of microRNA (miRNA) target sites, we provide evidence for preexisting heterogeneities in dosage sensitivity among genes on the mammalian X and avian Z chromosomes. We first showed that, across all human autosomal genes, dosage sensitivity -- as indicated by patterns of genic copy number variation -- correlates with the degree of conserved miRNA targeting. We found that conserved targeting correlates especially strongly with sensitivity to dosage increases, consistent with miRNA targeting serving to reduce gene expression. Turning to the sex chromosomes of mammals and birds, genes that retained a homolog on the sex-specific Y or W chromosome (X-Y and Z-W pairs) have more conserved miRNA target sites than genes with no Y or W homolog. In mammals, genes with no Y homolog that became subject to XCI have more conserved sites than those that continued to escape XCI following Y gene decay. In birds, across Z-linked genes with no W homolog, the degree of conserved miRNA targeting correlates with the degree of gene-specific dosage compensation. We then reconstructed the ancestral state of miRNA targeting, observing significant heterogeneities in the extent of miRNA targeting, and thus dosage sensitivity, on the ancestral autosomes that gave rise to the mammalian and avian sex chromosomes. Finally, through analysis of publically available experimental datasets, we validated the function, in living cells,of the miRNA target sites used to infer dosage sensitivity. We thus conclude that differences in dosage sensitivity – both to increases and to decreases in gene dosage -- among genes on the ancestral autosomes influenced their evolutionary trajectory during sex chromosome evolution, not only on the sex-specific Y and W chromosomes, but also on the sex-shared X and Z chromosomes.

Our findings build upon previous work in three important ways. First, our analysis of miRNA-mediated repression indicates that these heterogeneities consist of sensitivities to dosage increases and decreases, whereas previous studies had either focused on sensitivity to underexpression or could not differentiate the two. Second, our reconstruction of miRNA targeting on the ancestral autosomes provides direct evidence that heterogeneities in dosage sensitivity among classes of X- and Z-linked were preexisting rather than acquired during sex chromosome evolution. Finally, by pointing to specific regulatory sequences (miRNA target sites) functioning to tune gene dosage both prior to and during sex chromosome evolution, our study provides a view of dosage compensation encompassing post-transcriptional regulation.

Human disease studies support the claim that increased dosage of X-Y pairs and X-inactivated genes is deleterious to fitness. Copy number gains of the X-linked gene *KDM6A*, which has a surviving human Y homolog, are found in patients with developmental abnormalities and intellectual disability (Lindgren et al., 2013). *HDAC6*, *CACNA1F*, *GDI1*, and *IRS4* all lack Y homologs and are subject to XCI in humans. A mutation in the 3 ̀ UTR of *HDAC6* abolishing targeting by miR-433 has been linked to familial chondrodysplasia in both sexes (Simon et al., 2010). Likely gain-of-function mutations in *CACNA1F* cause congenital stationary night blindness in both sexes (Hemara-Wahanui et al., 2005). Copy number changes of *GDI1* correlate with the severity of X-linked mental retardation in males, with female carriers preferentially inactivating the mutant allele (Vandewalle et al., 2009). Somatic genomic deletions downstream of *IRS4* lead to its overexpression in lung squamous carcinoma (Weischenfeldt et al., 2017). Males with partial X disomy due to translocation of the distal long arm of the X chromosome (Xq28) to the long arm of the Y chromosome show severe mental retardation and developmental defects (Lahn et al., 1994). Most genes in Xq28 are inactivated in 46,XX females but escape inactivation in such X;Y translocations, suggesting that increased dosage of Xq28 genes caused the cognitive and developmental defects. We anticipate that further studies will reveal additional examples of the deleterious effects of increases in gene dosage of X-Y pairs and X-inactivated genes.

We and others previously proposed that Y gene decay drove upregulation of homologous X-linked genes in both males and females, and that XCI subsequently evolved at genes sensitive to increased expression from two active X-linked copies in females (Jegalian & Page, 1998; Ohno, 1967). Our finding that X-inactivated genes have higher miRNA conservation scores than X escape genes is consistent with this aspect of the model. However, recent studies indicating heterogeneity in dosage sensitivity between classes of mammalian X- or avian Z-linked genes (Bellott et al., 2014, 2017; Pessia et al., 2012; Zimmer et al., 2016), combined with the present finding that these dosage sensitivities existed on the ancestral autosomes, challenge the previous assumption of a single evolutionary pathway for all sex-linked genes.

We therefore propose a revised model of X-Y and Z-W evolution in which the ancestral autosomes that gave rise to the mammalian and avian sex chromosomes contained three (or two, in the case of birds) classes of genes with differing dosage sensitivities (Figure 7A,B). For ancestral genes with high dosage sensitivity, Y or W gene decay would have been highly deleterious, and thus the Y- or W-linked genes were retained. According to our model, these genes’ high dosage sensitivity also precluded upregulation of the X- or Z-linked homolog, and, in mammals, subsequent X-inactivation; indeed, their X-linked homologs continue to escape XCI (Bellott et al., 2014). For ancestral mammalian genes of intermediate dosage sensitivity, Y gene decay did occur, and was accompanied or followed by compensatory upregulation of the X-linked homolog in both sexes; the resultant increased expression in females was deleterious and led to the acquisition of XCI. Ancestral mammalian genes of low dosage sensitivity continued to escape XCI following Y decay; heterogeneity in X upregulation may further subdivide such genes (Figure 6A). These genes’ dosage insensitivity set them apart biologically, and evolutionarily, from the other class of X-linked genes escaping XCI -- those with a surviving Y homolog.

**Figure 7.**
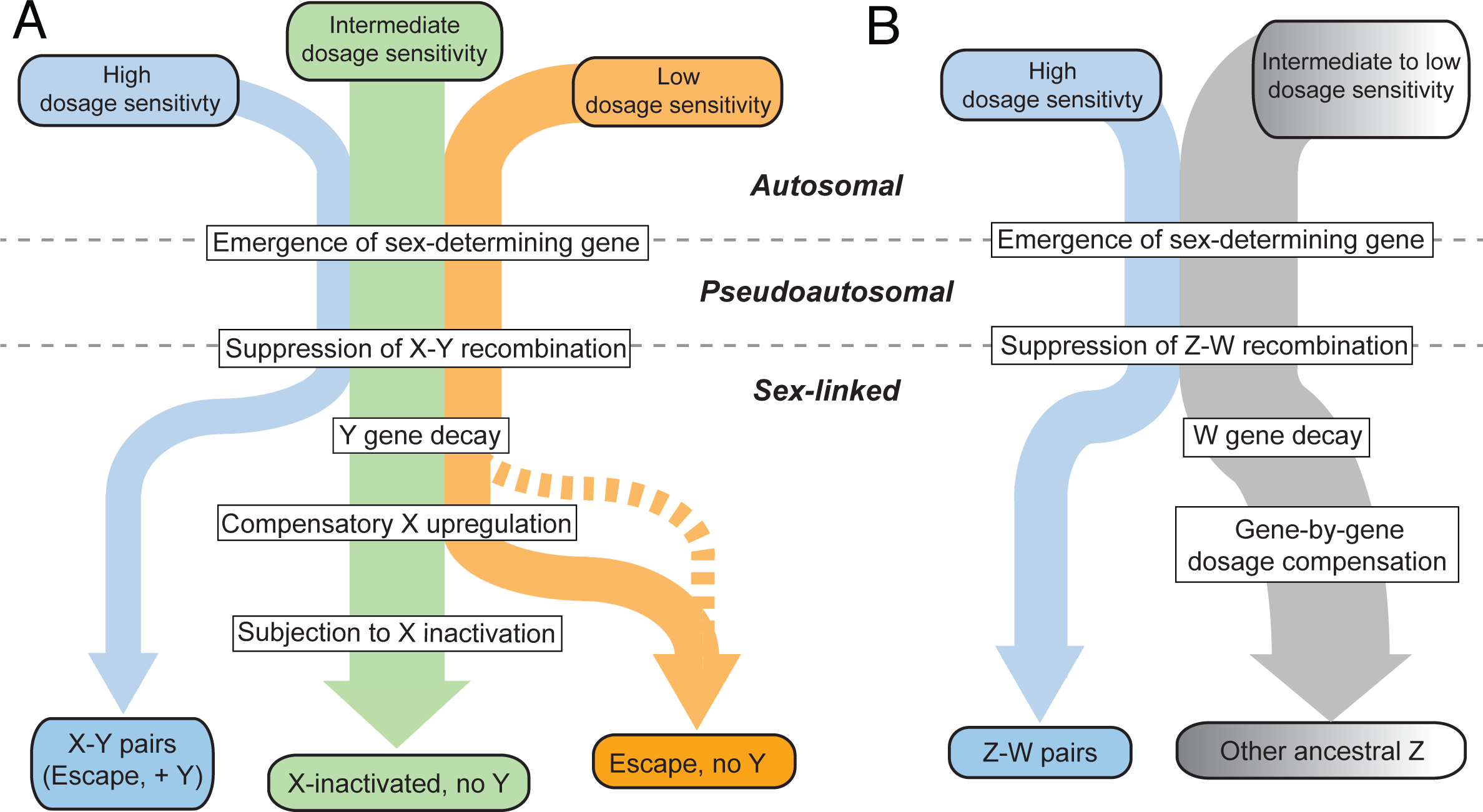
An evidence-based model of preexisting heterogeneities in dosage sensitivity shaping mammalian and avian sex chromosome evolution. In this model, preexisting heterogeneities in dosage sensitivity determined the trajectory of Y/W gene loss in both mammals and birds, and of subsequent X-inactivation in mammals and dosage compensation in birds. Colored arrow widths are scaled approximately to the number of ancestral genes in each class. (A) The dashed orange line represents the possibility that a subset of X-linked genes may have not undergone compensatory X upregulation following Y gene decay. (B) Ancestral Z genes with no W homolog follow a gradient of preexisting dosage sensitivity (top, grey to white), which determined the degree of dosage compensation following W gene loss (bottom).

Our revised model relates preexisting, gene-by-gene heterogeneities in dosage sensitivity to the outcomes of sex chromosome evolution. However, the suppression of X-Y recombination did not occur on a gene-by-gene basis, instead initiating Y gene decay and subsequent dosage compensation through a series of large-scale inversions encompassing many genes (Lahn & Page, 1999). The timings and boundaries of these evolutionary strata varied among mammalian lineages, thus leading to unique chromosome-scale evolutionary dynamics across mammals. These large-scale changes would have then allowed for genic selection to take place according to the preexisting dosage sensitivities outlined above. In this way, the course of sex chromosome evolution in mammals is a composite of 1) preexisting, gene-by-gene dosage sensitivities and 2) the manner in which the history of the X and Y unfolded in particular lineages via discrete, large-scale inversions.

In this study, we have focused on classes of ancestral X-linked genes delineated by the survival of a human Y homolog or by the acquisition of XCI in humans, but such evolutionary states can differ among mammalian lineages and species. In mouse, for instance, both Y gene decay (Bellott et al., 2014) and the acquisition of X-inactivation (Yang et al., 2010) are more complete than in humans or other mammals, as exemplified by *RPS4X*, which retains a Y homolog and continues to escape XCI in primates, but has lost its Y homolog and is subject to XCI in rodents. These observations could be explained by shortened generation times in the rodent lineage, resulting in longer evolutionary times, during which the forces leading to Y gene decay and the acquisition of X-inactivation could act (Charlesworth & Crow, 1978; Jegalian & Page, 1998; Ohno, 1967). Another case of lineage differences involves *HUWE1*, which lacks a Y homolog and is subject to XCI in both human and mouse, but retains a functional Y homolog in marsupials, where it continues to escape XCI. In the future, more complete catalogues of X-inactivation and escape in additional mammalian lineages would make it possible to examine whether analogous, preexisting dosage sensitivities differentiate the three classes of X-linked genes (X-Y pairs, X-inactivated genes, and X escape genes) in other species.

Previous studies have sought evidence of X-linked upregulation during mammalian sex chromosome evolution using comparisons of gene expression levels between the whole of the X chromosome and all of the autosomes, with equal numbers of studies rejecting or finding evidence consistent with upregulation (Deng et al., 2011; Julien et al., 2012; Kharchenko, Xi, & Park, 2011; Lin, Xing, Zhang, & He, 2012; Xiong et al., 2010). This is likely due to gene-by-gene heterogeneity in dosage sensitivities that resulted in a stronger signature of upregulation at more dosage sensitive genes (Pessia et al., 2012). Similarly, studies of Z-linked gene expression in birds provide evidence for the gene-by-gene nature of Z dosage compensation, as measured by comparisons of gene expression levels between ZZ males and ZW females (Itoh et al., 2007; Mank & Ellegren, 2009; Uebbing et al., 2015), and indicate a stronger signature of dosage compensation at predicted dosage-sensitive genes (Zimmer et al., 2016). By showing that such dosage sensitivities existed on the ancestral autosomes and consist of sensitivity to both increases and decreases, our findings highlight an additional aspect of dosage compensation that affects both birds and mammals.

In addition to revealing similarities between mammals and birds, our study provides a view of dosage compensation that highlights post-transcriptional regulatory mechanisms, pointing to specific non-coding sequences with known mechanisms (microRNA target sites) functioning across evolutionary time. A recent study in birds showed a role for a Z-linked miRNA, miR-2954-3p, in dosage compensation of some Z-linked genes (Warnefors et al., 2017). Our study suggests an additional, broader role for miRNA targeting, with hundreds of different miRNAs acting to tune gene dosage both before and during sex chromosome evolution. Furthermore, our finding of greater conserved miRNA targeting of X-inactivated genes relative to X escape genes shows that it is possible to predict the acquisition of a transcriptional regulatory state (XCI) during sex chromosome evolution on the basis of a preexisting, post-transcriptional regulatory state. Perhaps additional post-transcriptional regulatory mechanisms and their associated regulatory elements will be shown to play roles in mammalian and avian dosage compensation.

Recent work has revealed that the sex-specific chromosome -- the Y in mammals and the W in birds -- convergently retained dosage-sensitive genes with important regulatory functions (Bellott et al., 2014, 2017). Our study, by reconstructing the ancestral state of post-transcriptional regulation, provides direct evidence that such heterogeneity in dosage sensitivity existed on the ancestral autosomes that gave rise to the mammalian and avian sex chromosomes. This heterogeneity influenced both survival on the sex-specific chromosomes in mammals and birds and the evolution of XCI in mammals. Thus, two independent experiments of nature offer empirical evidence that modern-day amniote sex chromosomes were shaped, during evolution, by the properties of the ancestral autosomes from which they derive.

## METHODS

### Statistics

Details of all statistical tests (type of test, test statistic, and p-value) used in this manuscript are provided in Supplemental Table S1.

### Human genic copy number variation

To annotate gene deletions and duplications, we used data from the Exome Aggregation Consortium (ExAC) (ftp://ftp.broadinstitute.org/pub/ExAC_release/release0.3.1/cnv/), which consists of autosomal genic duplications and deletions (both full and partial) called in 59,898 exomes (Ruderfer et al., 2016). Further details are provided in Supplemental Methods in the section titled ‘Human genic copy number variation’. These gene assignments are provided in Supplemental Table S2.

### X- and Z-linked gene sets

We utilized our previous reconstructions of the ancestral mammalian X (Bellott et al., 2014) and avian Z (Bellott et al., 2017) chromosomes, as well as information on multicopy and ampliconic X-linked genes (Mueller et al., 2013) and XCI status in humans (Balaton et al., 2015) to delineate classes of X- and Z-linked genes. Further details are provided in Supplemental Methods under the sections titled ‘X-linked gene sets’ and ‘Z-linked gene sets’. Information on X-linked genes is provided in Supplemental Table S3. Information on Z-linked genes is provided in Supplemental Table S4.

### microRNA target sites

Pre-calculated P_CT_ scores for all gene-miRNA family interactions

(http://www.targetscan.org/vert71/vert71datadownload/SummaryCounts.allpredictions.txt.zip) and site-wise alignment information

(http://www.targetscan.org/vert71/vert71datadownload/ConservedFamilyInfo.txt.zip) were obtained from TargetScan Human v7. Details on the filtering of miRNAs and resampling-based assessment of P_CT_ scores are provided in Supplemental Methods in the section titled ‘microRNA target site P_CT_ scores’. Details regarding analysis of human-chicken or human-anolis conserved sites, as well as approaches to control for background conservation, are provided in Supplemental Methods in the section titled ‘Human-chicken conserved microRNA target sites.’

### Variation in within-UTR conservation bias

To address the possibility that non-uniformity in regional 3 ̀ UTR conservation could artificially inflate or deflate conservation scores of certain target sites, we implemented a step-detection algorithm to segment 3 ̀ UTRs into regions of homogeneous background conservation and calculated miRNA site conservation relative to these smaller regions. These regionally normalized scores, corresponding to all gene-miRNA interactions, are provided in Supplemental Table S5. Details of the step-detection algorithm are provided in Supplemental Methods in the section titled ‘Variation in within-UTR conservation bias’.

### Logistic regression

Logistic regression models were constructed using the function ‘multinom’ in the R package ‘nnet.’ We used previously published values for known factors in the survival of Y-linked (Bellott et al., 2014) and W-linked (Bellott et al., 2017) genes except for human expression breadth, which we recalculated using data from the GTEx Consortium v6 data release (Consortium, 2015). Briefly, kallisto was used to estimate transcript per million (TPM) values in the 10 male samples with the highest RNA integrity numbers (RINs) from each of 37 tissues, and expression breadth across tissues was calculated as described in (Bellott et al., 2014), using median TPM values for each tissue.

### Assessing Z-linked dosage compensation using cross-species RNA-sequencing data

Raw data from Marin et al 2017 (add citation) was obtained, and kallisto and limma/voom were used for abundance quantification and differential expression, respectively. Further details are provided in Supplemental Methods in the section titled ‘Assessing Z-linked dosage compensation using cross-species RNA-sequencing data’.

### Gene expression profiling and crosslinking datasets

Fold-changes in mRNA expression and targets of Argonaute as determined by high-throughput crosslinking-immunoprecipitation (CLIP) were obtained from a variety of publically available datasets. Further details are provided in Supplemental Methods in the section titled ‘Gene expression profiling and crosslinking datasets.’ All fold-changes and CLIP targets are provided in Supplemental Table S6.

### Code availability

A custom Python (RRID:SCR_008394) script utilizing Biopython (RRID:SCR_007173) was used to generate shuffled miRNA family seed sequences. Identification of miRNA target site matches using shuffled seed sequences was performed using the ‘targetscan_70.pl’ perl script (http://www.targetscan.org/vert71/vert71datadownload/targetscan70.zip). 3 ̀ UTR segmentation was performed with the ‘plot_transitions.py’ python script. Code is available at: https://github.com/snaqvi1990/Naqvi17-code and as Supplementary Code.

### Data access

Data supporting the findings of this study are available within the paper and its Supplemental information files. Accession numbers for publically available datasets are provided when appropriate in Methods sections.

## ACKNOWLEDGEMENTS

We thank V. Agarwal, S. Eichorn, S. McGeary, and D. Bartel for assistance with the TargetScan database and helpful discussions; A. Godfrey for updated human-chicken orthology information; and A. Godfrey, J. Hughes and H. Skaletsky for critical reading of the manuscript. This work was supported by the National Institutes of Health and the Howard Hughes Medical Institute. S.N. was supported under a research grant by Biogen.

## AUTHOR CONTRIBUTIONS

S.N., D.W.B. and D.C.P designed the study. S.N. performed analyses with assistance from D.W.B. K.S.L developed and implemented the step-detection algorithm. S.N. and D.C.P wrote the paper.

---

**Supplemental Figure S1:**
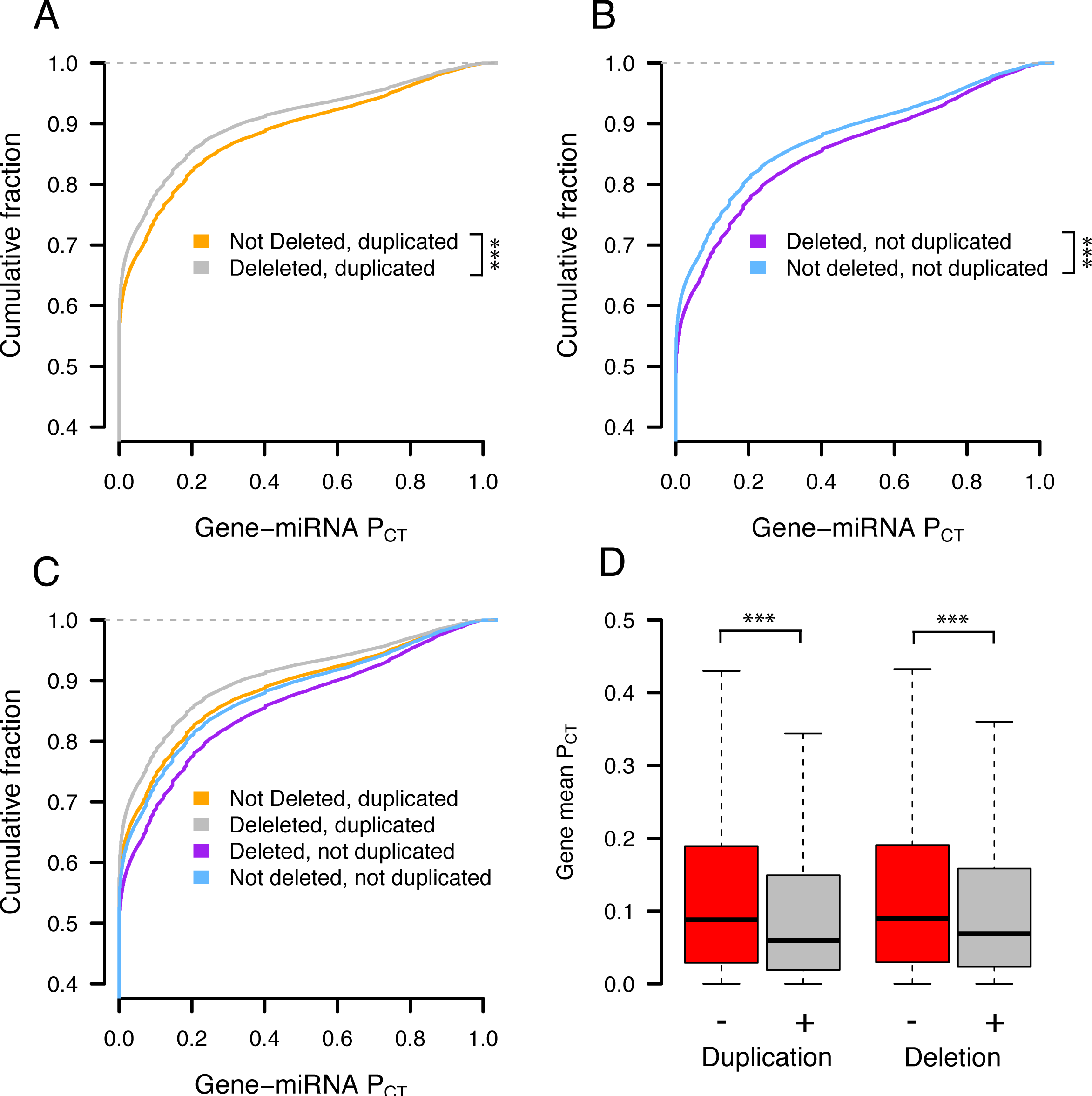
Effect of deletion status on autosomal P_CT_ scores. Probabilities of conserved targeting (P_CT_) of all gene-miRNA interactions involving non-deleted and deleted genes, further stratified as (A) duplicated (grey, n = 69,339 interactions from 4,118 genes; orange, n = 51,514 interactions from 2,916 genes) or (B) not duplicated (purple, n = 72,826 interactions from 3,510 genes; blue, n = 80,290 interactions from 3,976 genes). *** p < 0.001, two-sided Kolmogorov-Smirnov test. (C) P_CT_ scores for all gene sets in (A) and (B) superimposed on one plot. (D) Mean gene-level P_CT_ scores when aggregating sets of duplicated/not duplicated (left) or deleted/not deleted (right) genes. *** p < 0.0001, two-sided Wilcoxon rank-sum test.

**Supplemental Figure S2:**
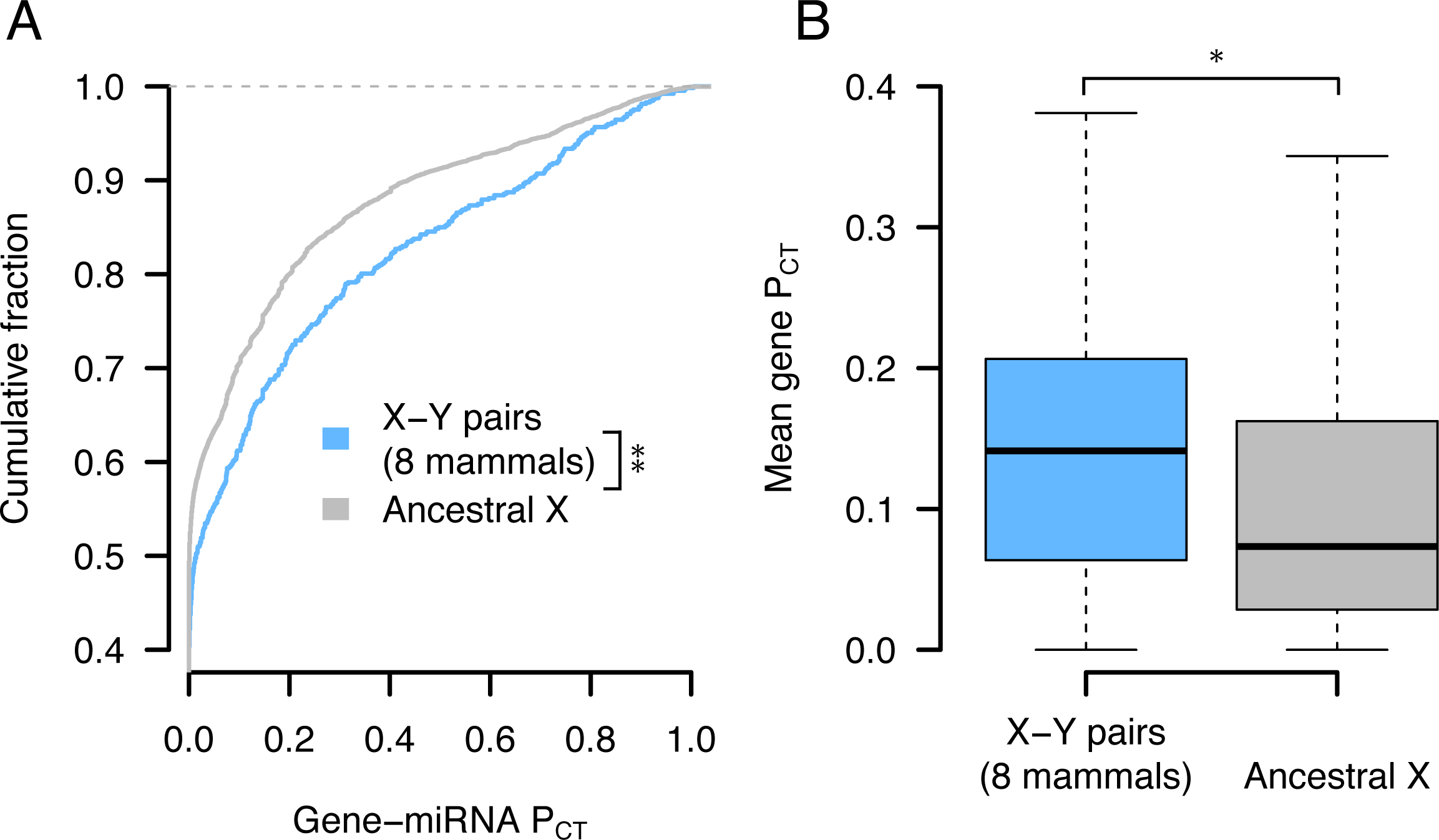
P_CT_ scores of X-Y pairs across 8 mammals. (A) P_CT_ score distributions of all gene-miRNA interactions involving X-Y pairs across eight sequenced mammalian Y chromosomes (n = 647 interactions from 32 genes) and other ancestral X genes (n = 8,831 interactions from 457 genes). ** p < 0.01, two-sided Kolmogorov-Smirnov test. (B) Gene-level mean P_CT_ scores. * p < 0.05, two-sided Wilcoxon rank-sum test.

**Supplemental Figure S3:**
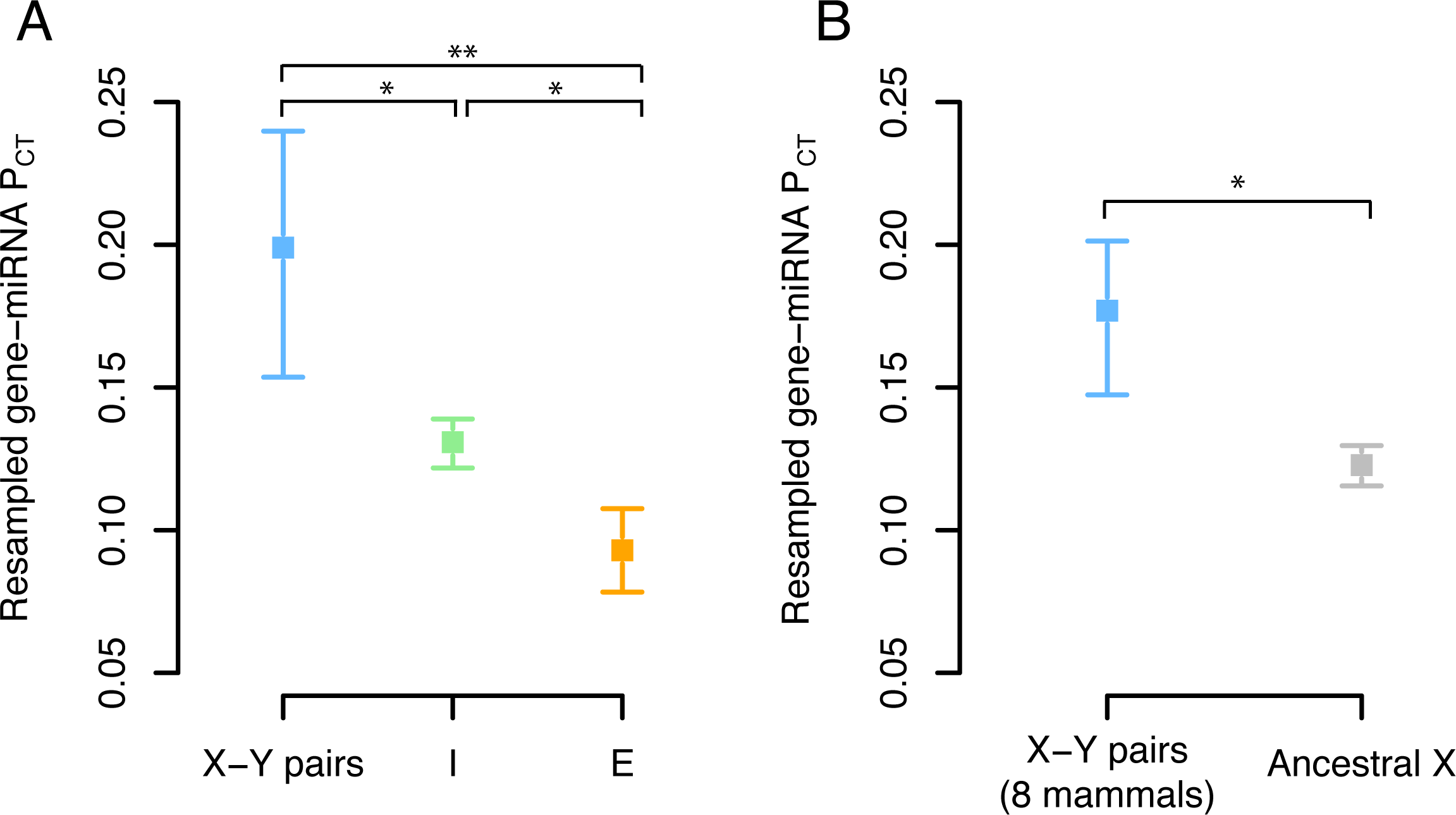
Resampled mean P_CT_ scores of X-linked genes. (A) Resampled gene-miRNA P_CT_ scores for human X-Y pairs (n = 15 genes), X-inactivated genes (n = 329 genes) and X escape genes (n = 56 genes). (B) Resampled gene-miRNA P_CT_ scores for X-Y pairs across eight mammals (n = 32 genes) and genes with no Y homolog in any of eight mammals (n = 457 genes). Points and error bars represent the median and 95% confidence intervals from 1,000 gene samplings with replacement. * p < 0.05, ** p < 0.01, empirical p-value computed as the fraction of random non-overlapping gene sets with a median difference in P_CT_ score at least as large as the true difference.

**Supplemental Figure S4:**
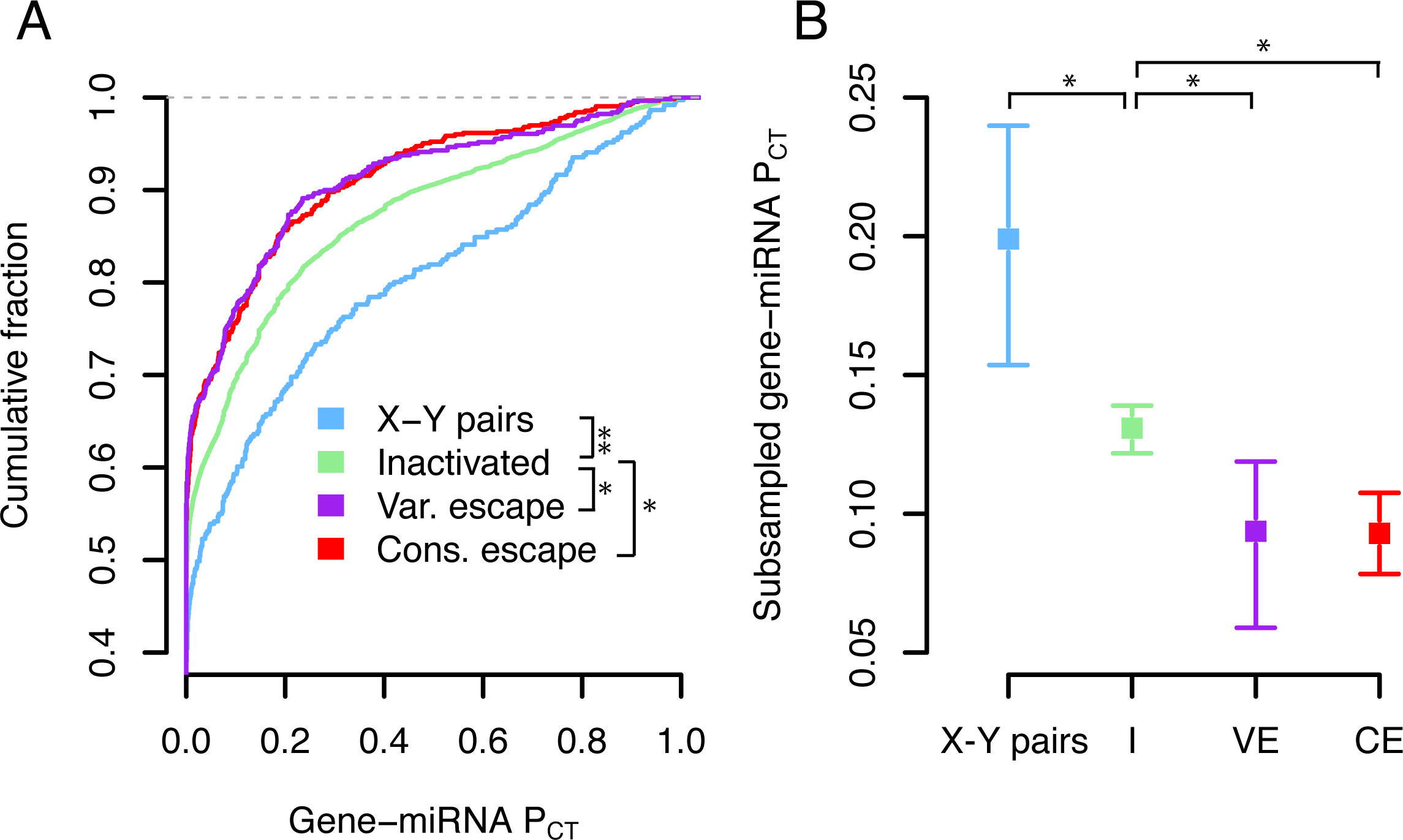
P_CT_ score comparisons with consistent and variable escape genes separated. (A) P_CT_ score distributions of all gene-miRNA interactions involving X-Y pairs (n = 371 interactions from 16 genes), X-inactivated genes (n = 6743 interactions from 329 genes), consistent escape genes (n = 567 interactions from 30 genes), or variable escape genes (n = 470 interactions from 26 genes) as defined by Balaton et al (Balaton et al., 2015). * p < 0.05, ** p < 0.01, two-sided Kolmogorov-Smirnov test. (B) Resampled gene-miRNA P_CT_ scores of gene classes from (A). Points and error bars represent the median and 95% confidence intervals from 1,000 gene samplings with replacement. * p < 0.05, empirical p-value computed as the fraction of random non-overlapping gene sets with a median difference in P_CT_ score at least as large as the true difference.

**Supplemental Figure S5:**
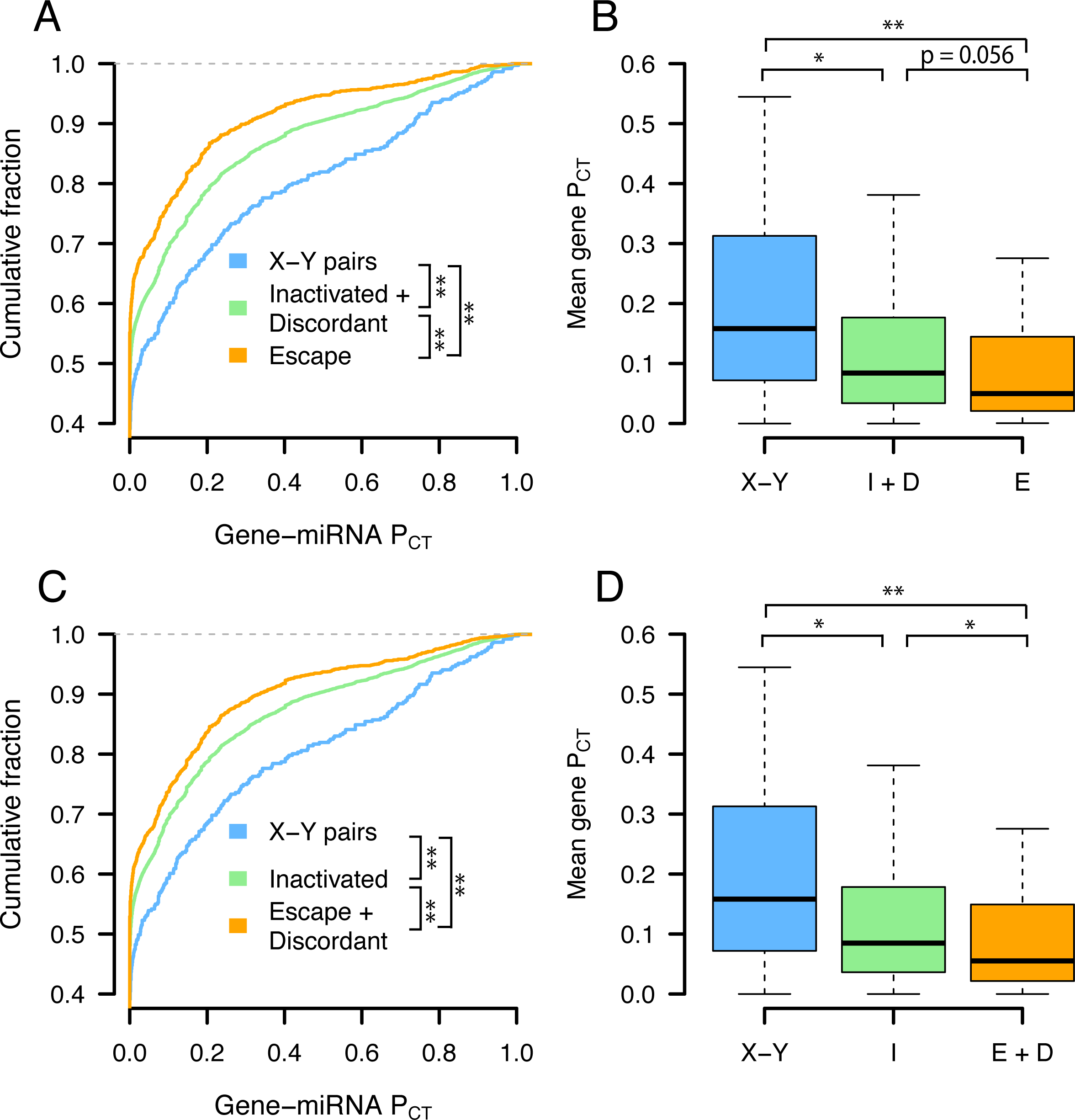
P_CT_ score comparisons with discordant genes included as X-inactivated or escape. P_CT_ score distributions of all gene-miRNA interactions (A,C) or mean gene-level P_CT_ score (B,D) of classes of X-linked genes with genes with a discordant XCI call (n = 721 interactions from 40 genes) included as X-inactivated (A,B) or X escape (C,D). Numbers of gene-miRNA interactions and genes as in Figure 1, but with the addition of discordant gene numbers/interactions to X-inactivated genes (A,B) or X escape genes (C,D). * p < 0.05, ** p < 0.01, two-sided Kolmogorov-Smirnov (A,C) or Wilcoxon rank-sum (B,D) test.

**Supplemental Figure S6:**
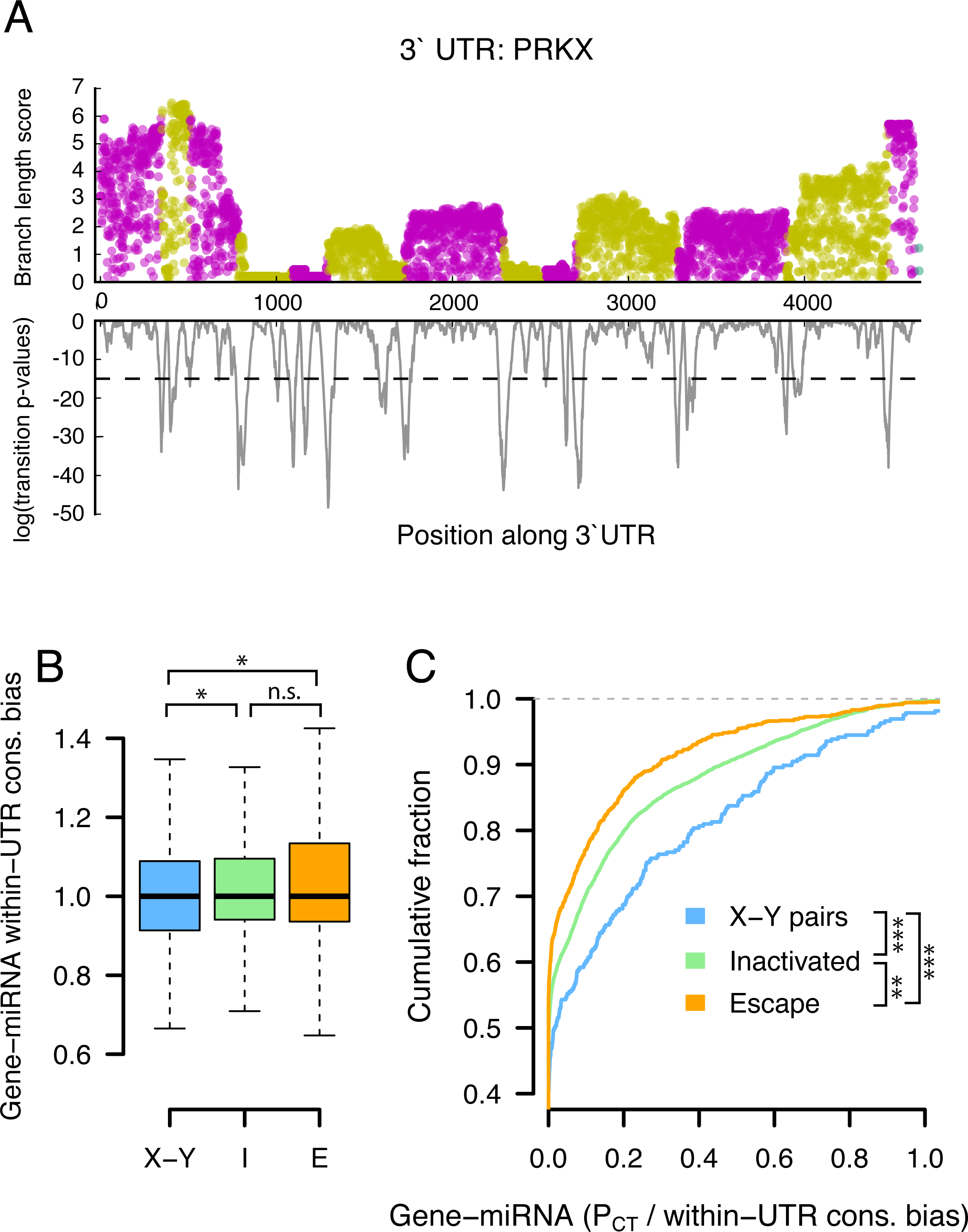
Variation in within-UTR conservation does not account for observed differences in P_CT_ score among classes of X-linked genes. (A) Example of step-detection to segment 3 ̀ UTRs. Top, base-wise branch length scores; bottom, probabilities of transition to a new section. Dashed line indicates p-value cutoff used to delineate a new section (plotted as alternating magenta/yellow points). (B) Boxplots of within-UTR conservation bias (see Methods) for all gene-miRNA interactions involving classes of X-linked genes. (C) Comparisons of P_CT_ scores normalized by within-UTR bias. **, p < 0.01, *** p < 0.001, two-sided Kolmogorov-Smirnov test.

**Supplemental Figure S7:**
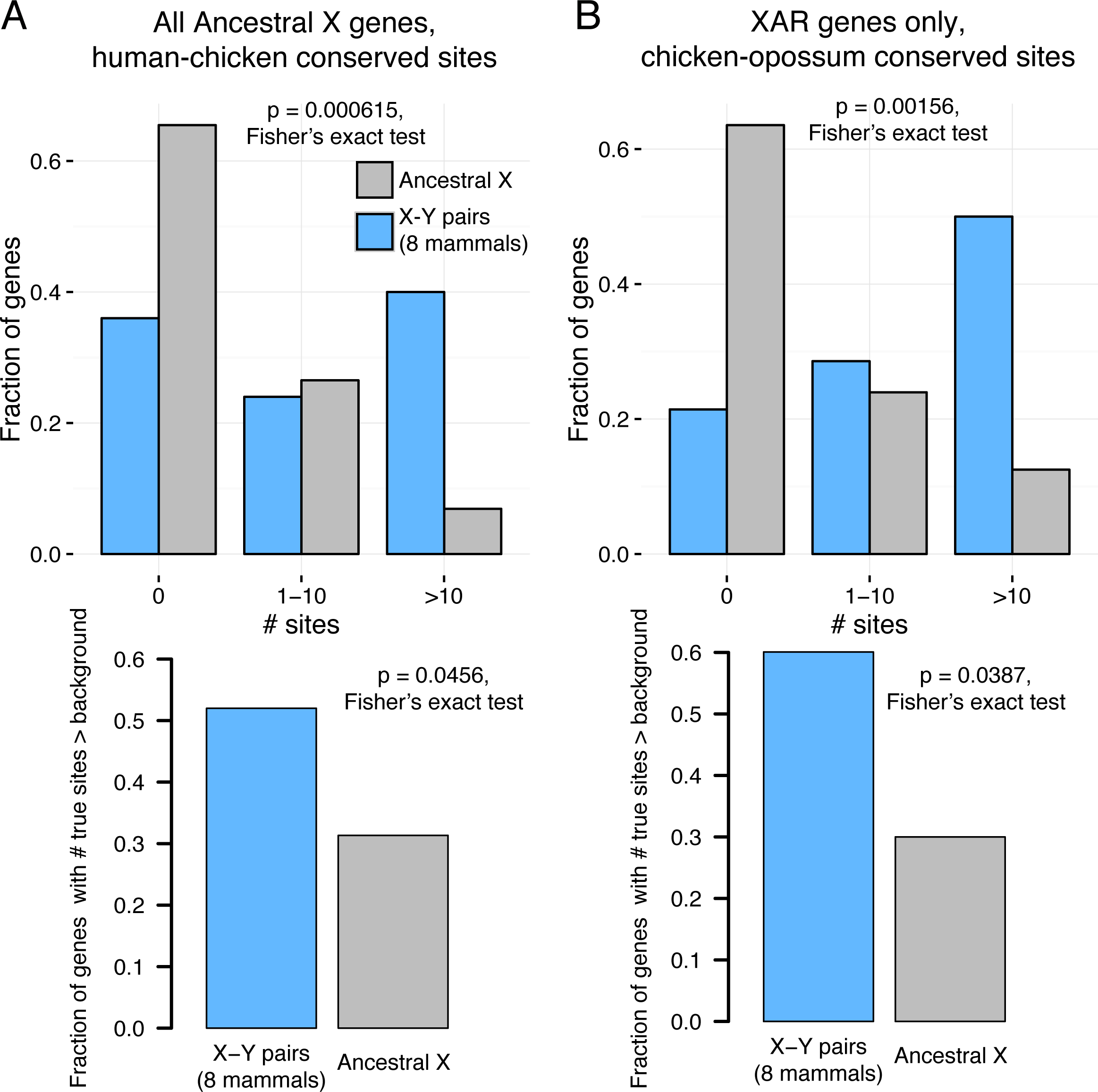
Ancestral miRNA targeting of X-Y pairs across 8 mammals. (A) Distributions of sites conserved between 3 ̀ UTRs of human and chicken orthologs (top) or comparisons to background expectation (bottom, see Methods) for X-Y pairs across 8 mammals (n = 25) and other ancestral X genes (n = 351). (D) Statistics as in (C), but using sites conserved between chicken and opossum 3 ̀ UTRs only for genes in the XAR; X-Y pairs across 8 mammals (n = 15), other ancestral X genes (n = 102).

**Supplemental Figure S8:**
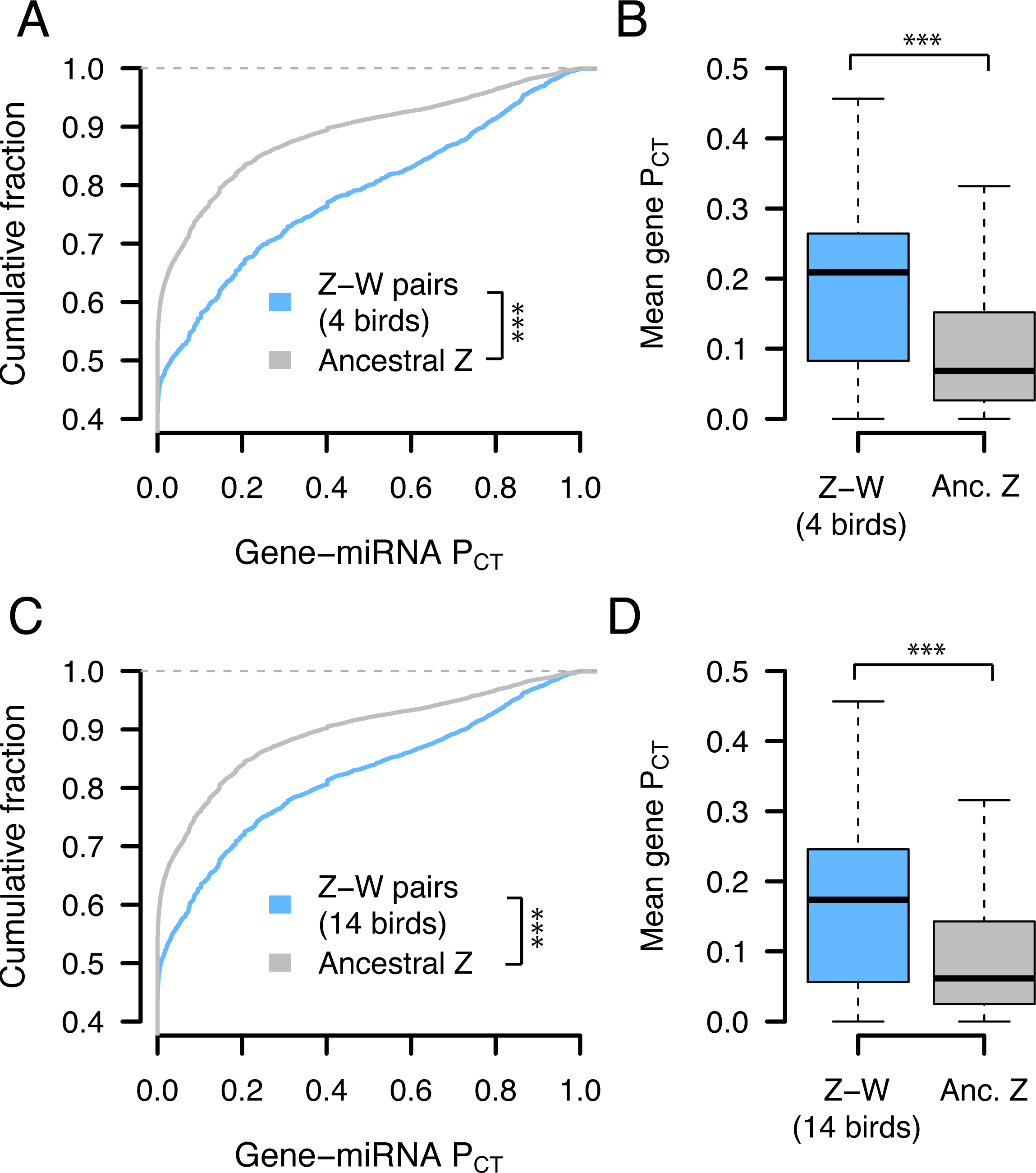
P_CT_ scores of Z-W pairs across 4 and 14 birds. (A,C) P_CT_ score distributions of all gene-miRNA interactions (A) Z-W pairs including predictions from three additional birds with male and female genome sequence (n = 2,187 interactions from 78 genes) and other ancestral Z genes (n = 15,357 interactions from 607 genes), or (C) Z-W pairs including read depth-based predictions from 10 additional birds with only female genome sequence (n = 4,458 interactions from 157 genes) and other ancestral Z genes (n = 13,086 interactions from 528 genes) *** p < 0.001, two-sided Kolmogorov-Smirnov test. (B,D) Gene-level mean p _CT_ scores. *** p < 0.01, two-sided Wilcoxon rank-sum test.

**Supplemental Figure S9:**
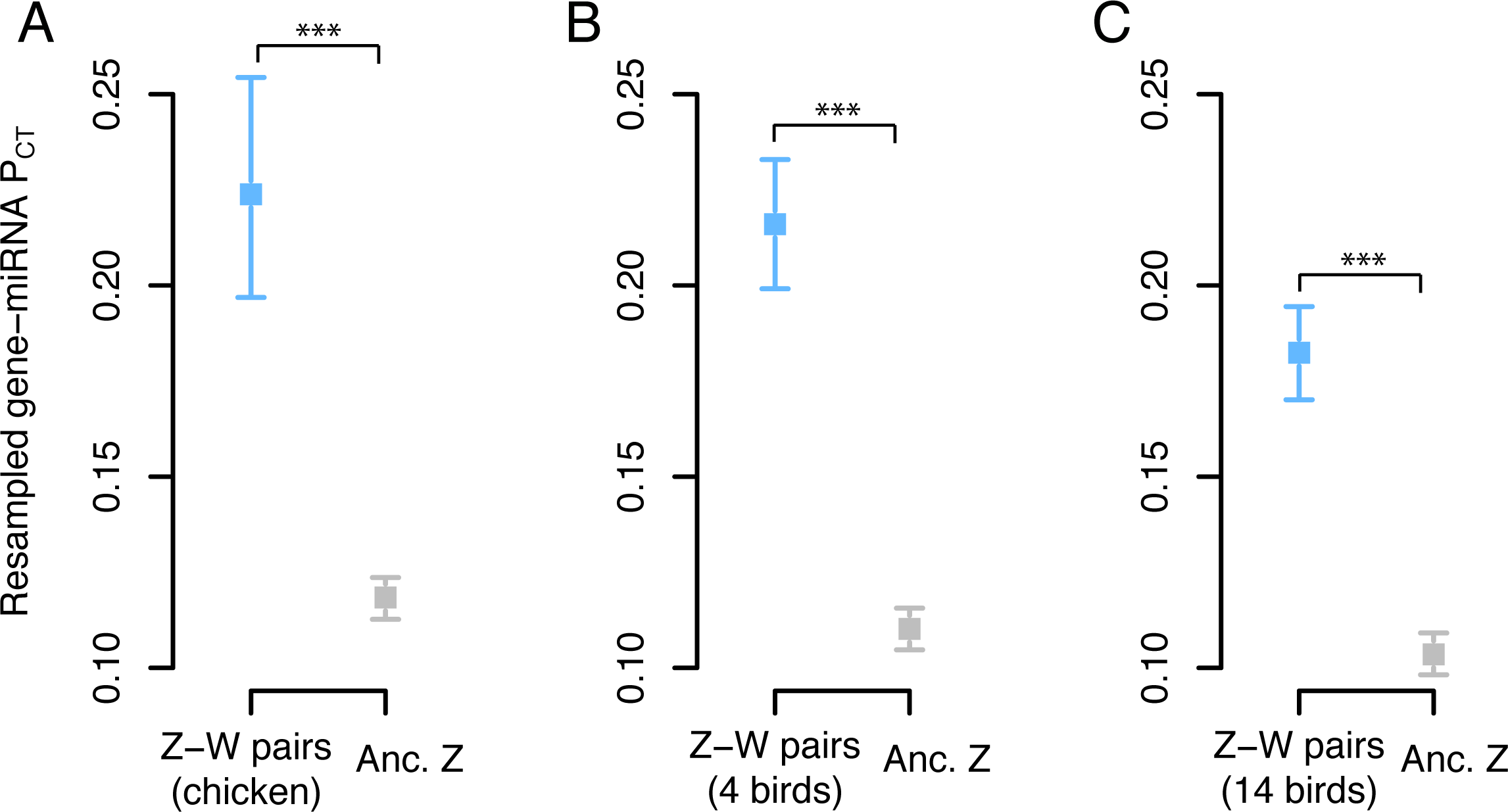
Resampled mean P_CT_ scores of Z-linked genes. Gene sets: (A) chicken Z-W pairs (n = 28 genes) and other ancestral Z genes (n = 657 genes), (B) Z-W pairs across four birds (n = 78 genes) compared to the remainder of ancestral Z genes (n = 607 genes), and (C) Z-W pairs across 14 birds (n = 157 genes) compared to the remainder of ancestral Z genes (n = 528 genes). Points and error bars represent the median and 95% confidence intervals from 1,000 gene samplings with replacement. *** p < 0.001, empirical p-value computed as the fraction of random non-overlapping gene sets with a median difference in P_CT_ score at least as large as the true difference.

**Supplemental Figure S10:**
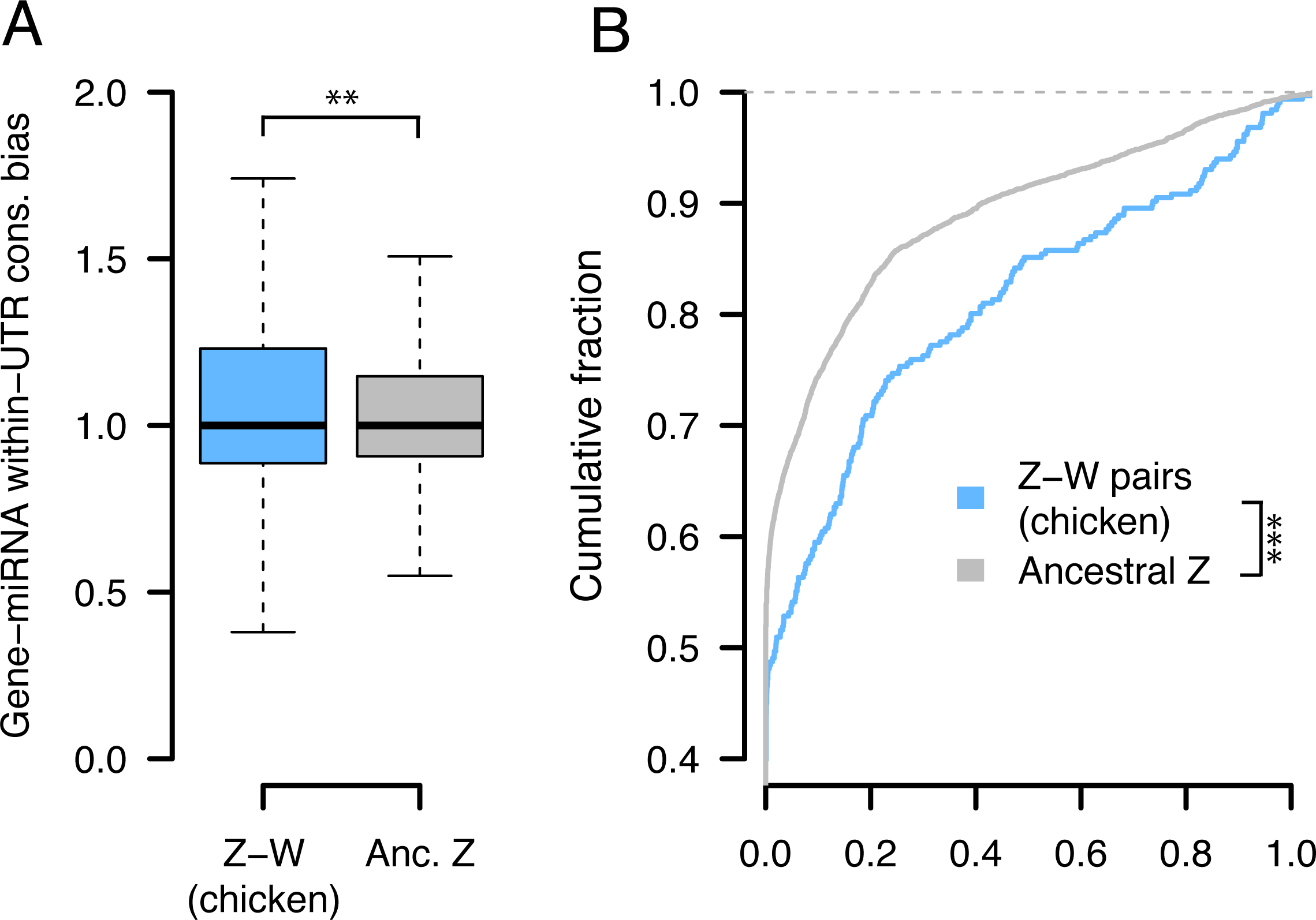
Variation in within-UTR conservation cannot account for observed differences in P_CT_ score among classes of Z-linked genes. (A) Boxplots of within-UTR conservation bias (see Methods) for all gene-miRNA interactions involving chicken Z-W pairs or other ancestral X genes. Numbers of interactions and genes as in Figure 4A. ** p < 0.01, two-side Wilcoxon rank-sum test. (B) Comparisons of P_CT_ scores normalized by within-UTR bias. *** p < 0.001, two-sided Kolmogorov-Smirnov test.

**Supplemental Figure S11:**
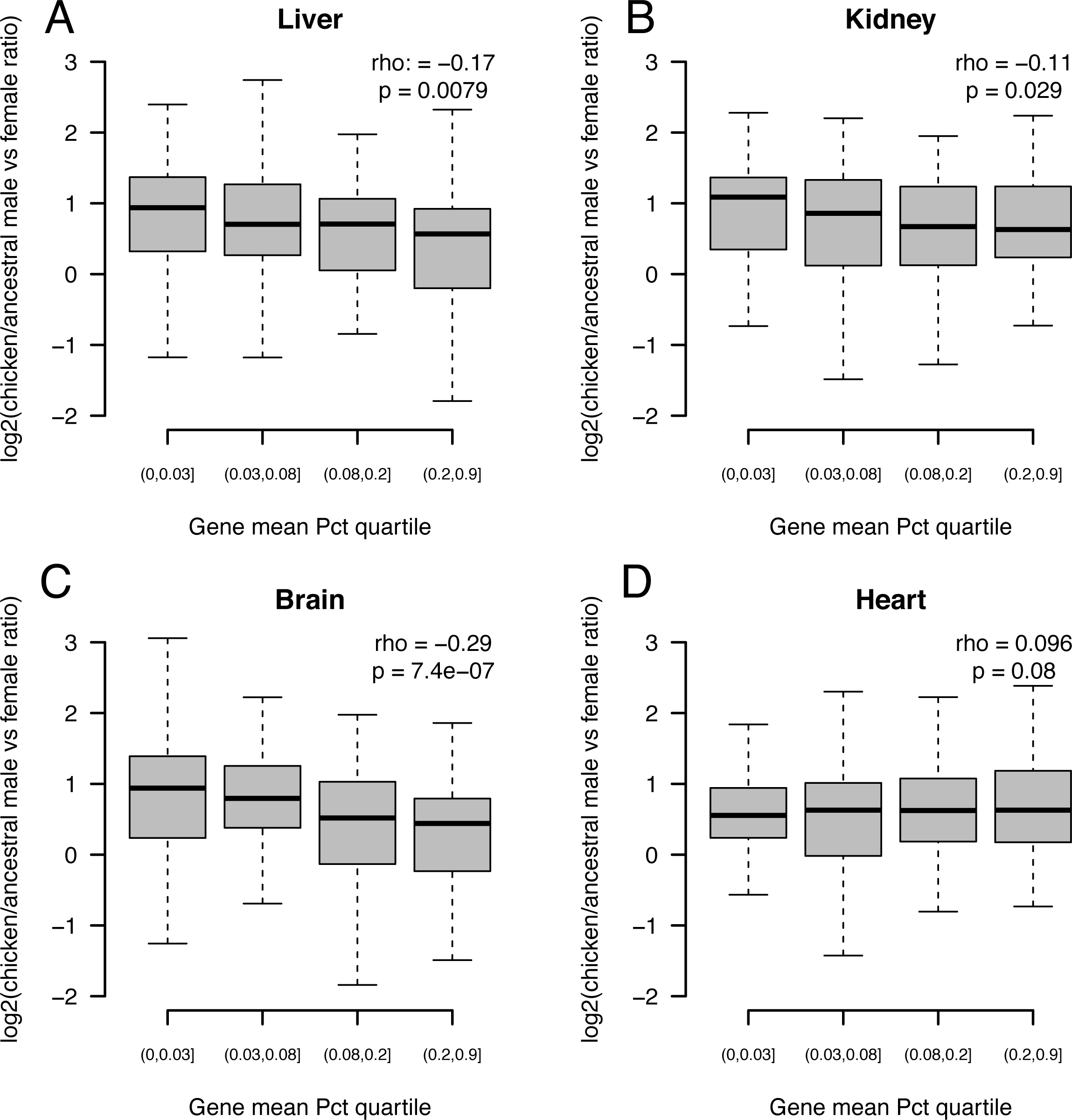
Correlation of Z-linked gene-specific dosage compensation with gene-level P_CT_ score. Distributions of chicken male/female expression ratio, normalized to that of human and anolis (y-axis) as a function of mean gene-level P_CT_ quartile (x-axis) for all expressed Z-linked gene with no W homolog. Expression ratios are plotted on a log_2_ scale; values closer to 0 imply more effective dosage compensation following W gene loss.

**Supplemental Figure S12:**
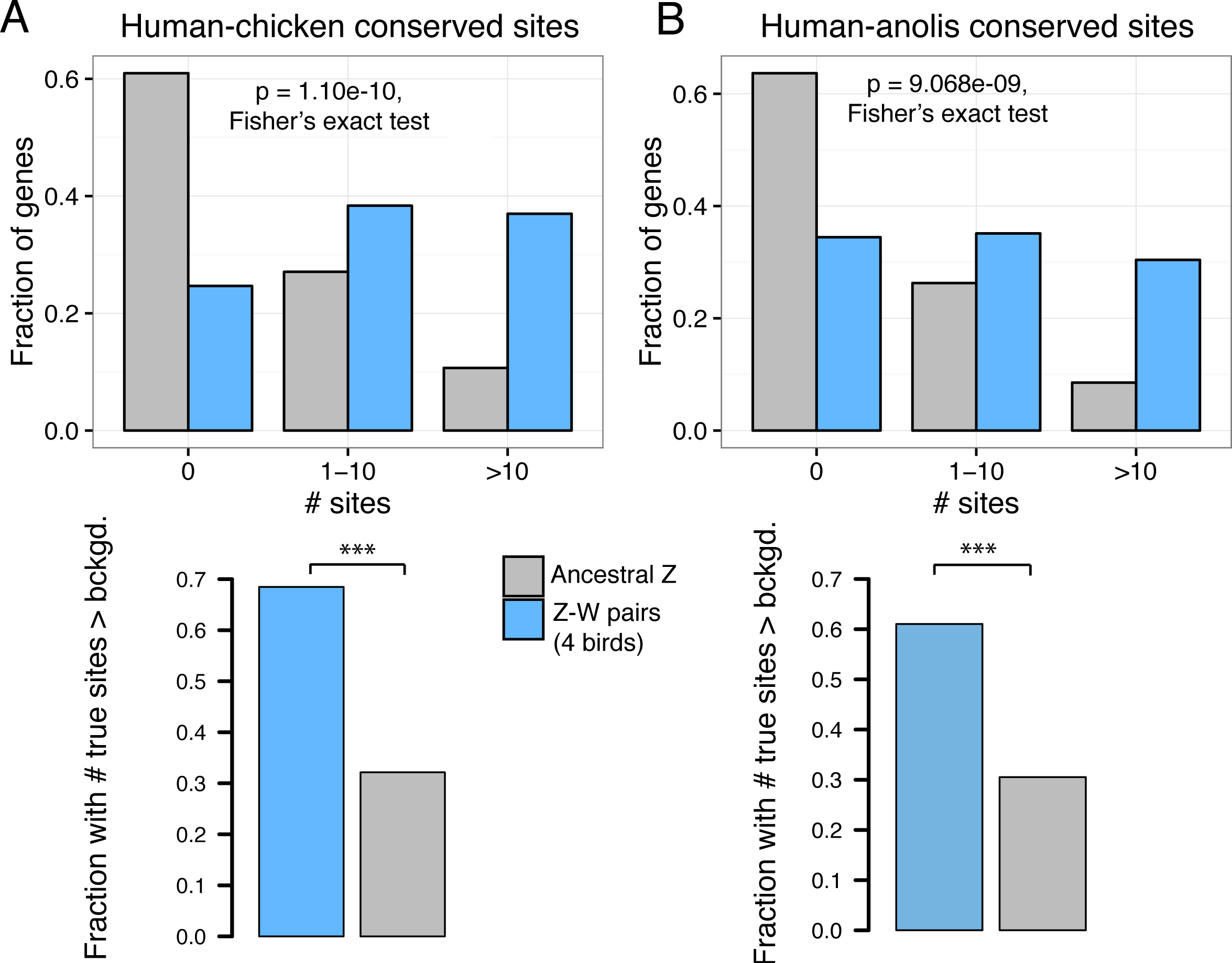
Ancestral miRNA targeting of Z-W pairs across 4 birds. (A) Distributions of sites conserved between 3 ̀ UTRs of human and chicken orthologs (top) or comparisons to background expectation (bottom, see Methods) for Z-W pairs across chicken and three additional birds with male and female genome sequence (4 birds, n = 73) and other ancestral Z genes (n = 532). (D) Statistics as in (C), but using sites conserved between human and anolis 3 ̀ UTRs; Z-W pairs across 4 birds (n = 73), other ancestral Z genes (n = 527). *** p < 0.001, two-sided Fisher’s exact test.

**Supplemental Figure S13:**
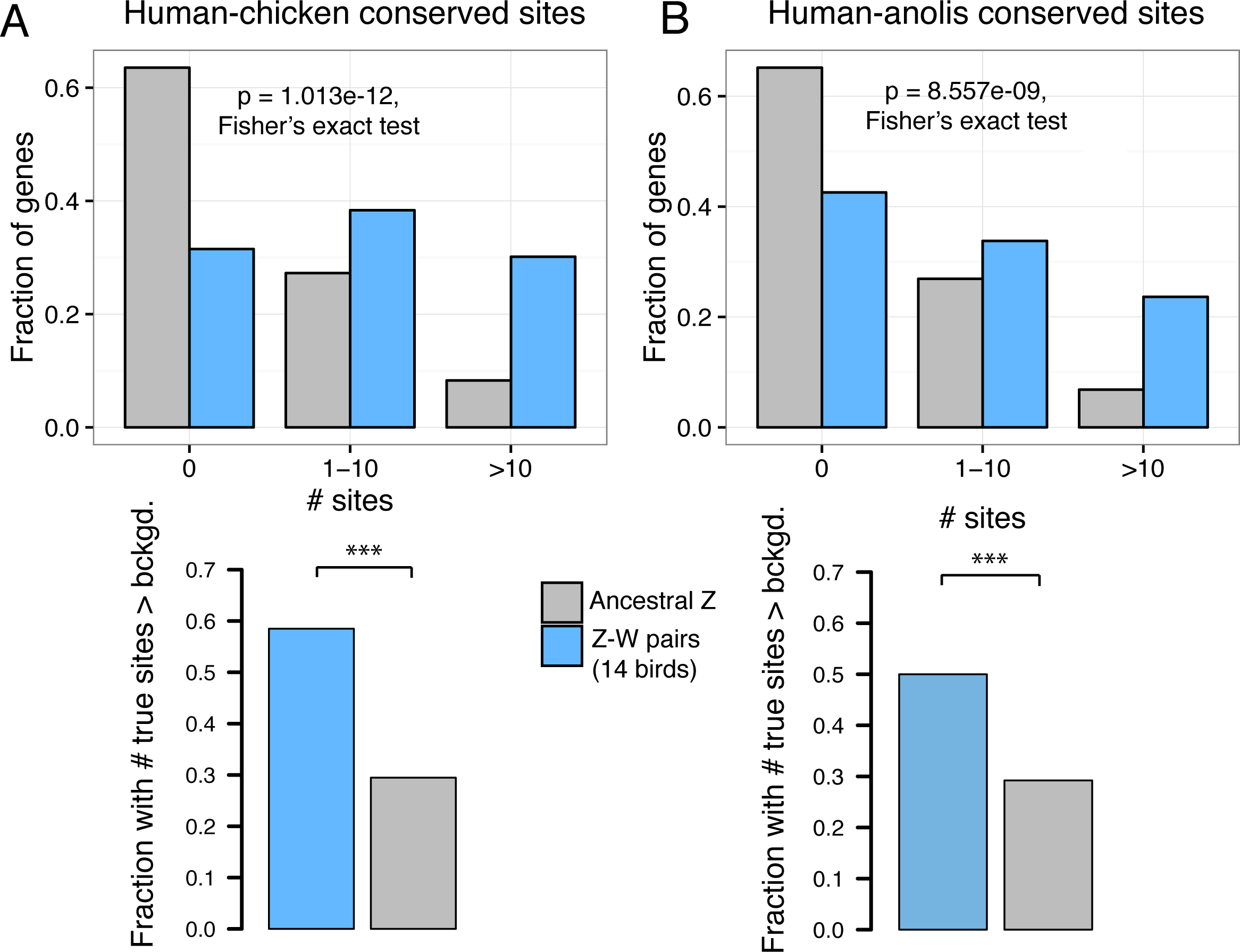
Ancestral miRNA targeting of predicted Z-W pairs across 14 birds. (A) Distributions of sites conserved between 3 ̀ UTRs of human and chicken orthologs (top) or comparisons to background expectation (bottom, see Methods) for Z-W pairs in chicken, predicted in three additional birds with male and female genome sequence, and predicted based on read depth from 10 additional birds with only female genome sequence (14 birds, n = 147) and other ancestral Z genes (n = 458). (D) Statistics as in (C), but using sites conserved between human and anolis 3 ̀ UTRs; Z-W pairs across 14 birds (n = 147), other ancestral Z genes (n = 453).

**Supplemental Figure S14:**
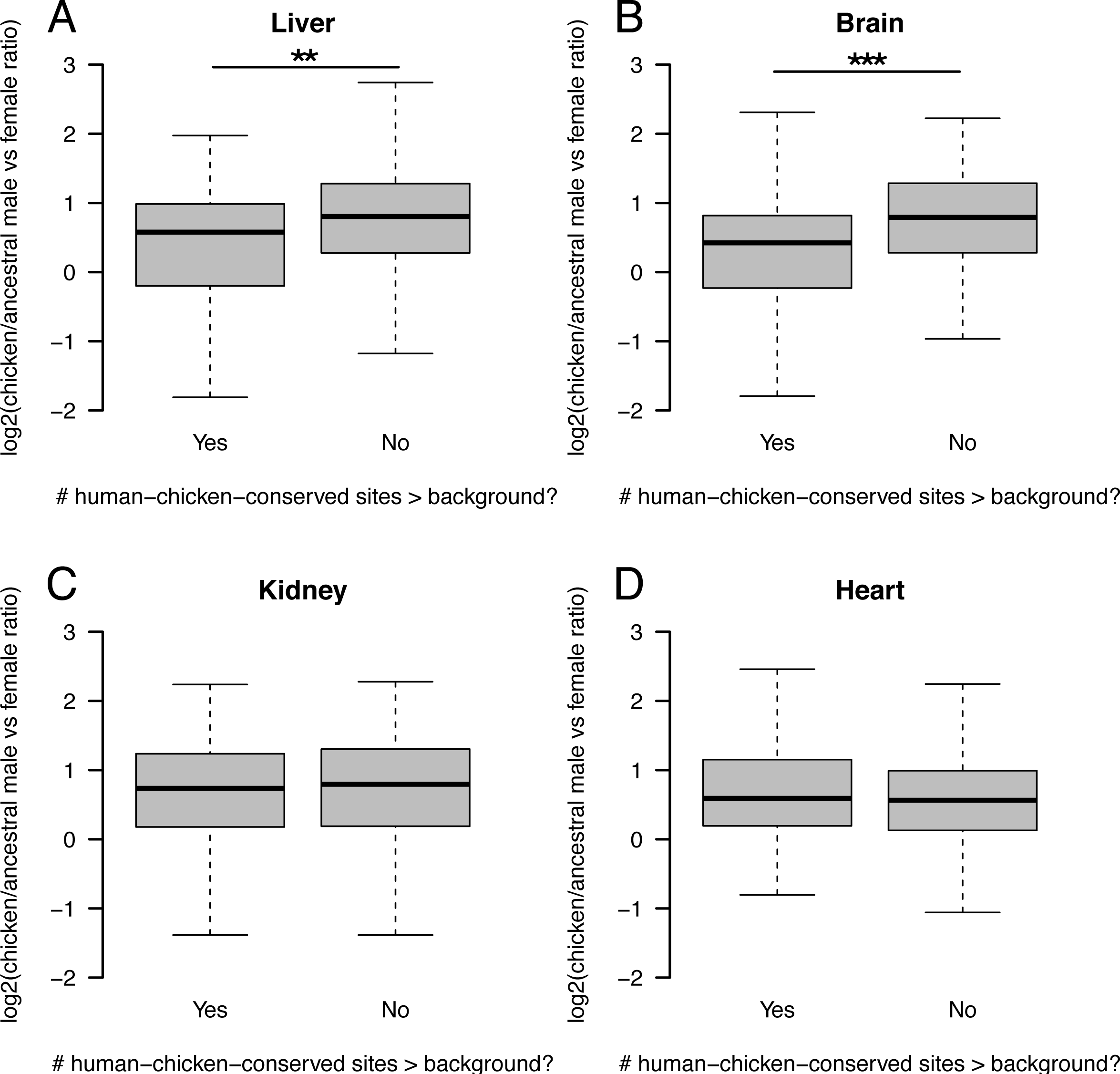
Correlation of Z-linked gene-specific dosage compensation with human-chicken-conserved site excess. Distributions of chicken male/female expression ratios, normalized to those of human and anolis (y-axis) for expressed Z-linked genes with no W homolog with (left) or without (right) an excess of human-chicken-conserved miRNA sites. ** p < 0.01, *** p < 0.0001, Wilcoxon rank-sum test.

**Supplemental Figure S15:**
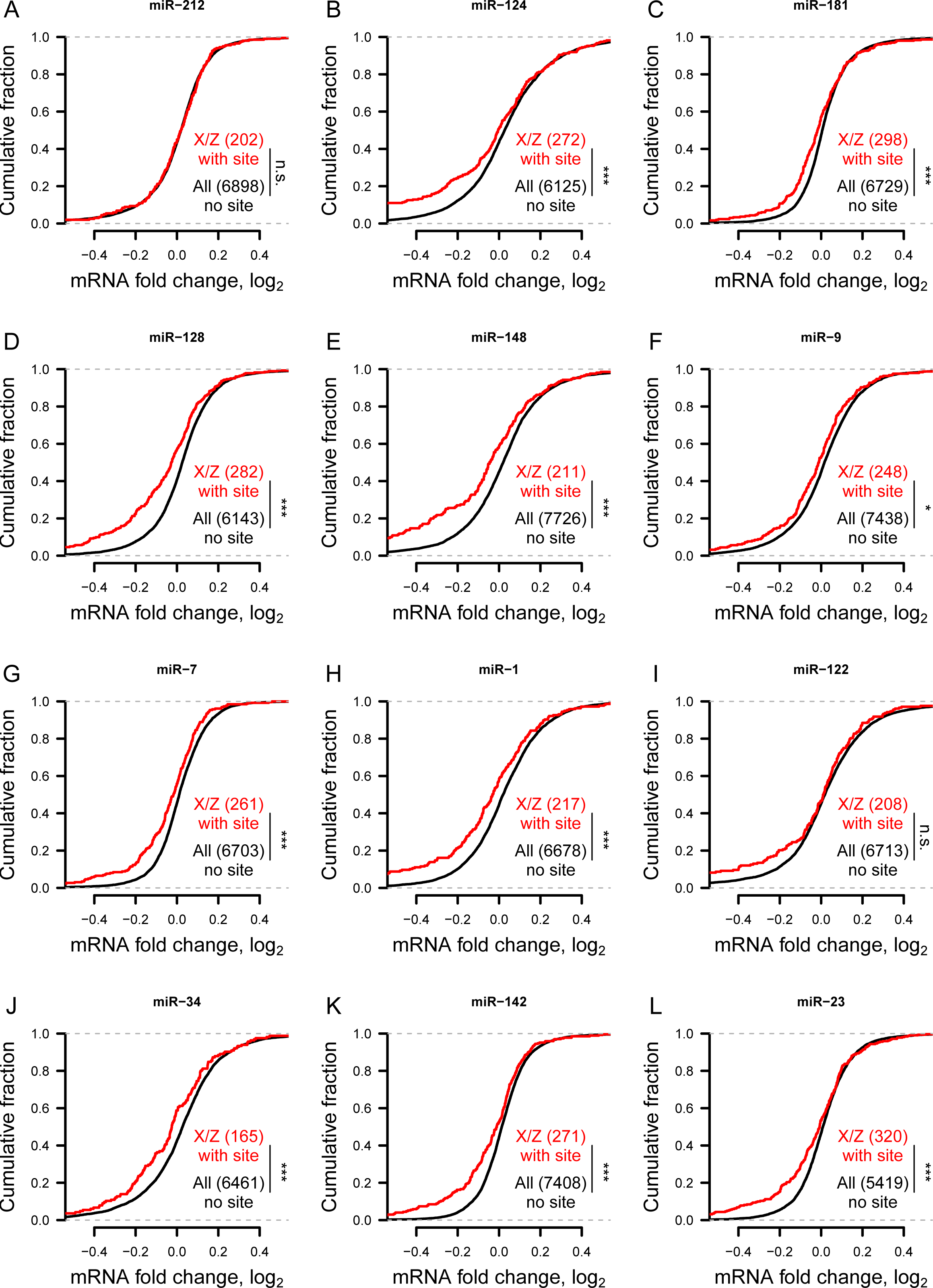
Gene expression changes following small RNA transfections in human HeLa cells. * p < 0.05, *** p < 0.001, two-sided K-S test.

**Supplemental Figure S16:**
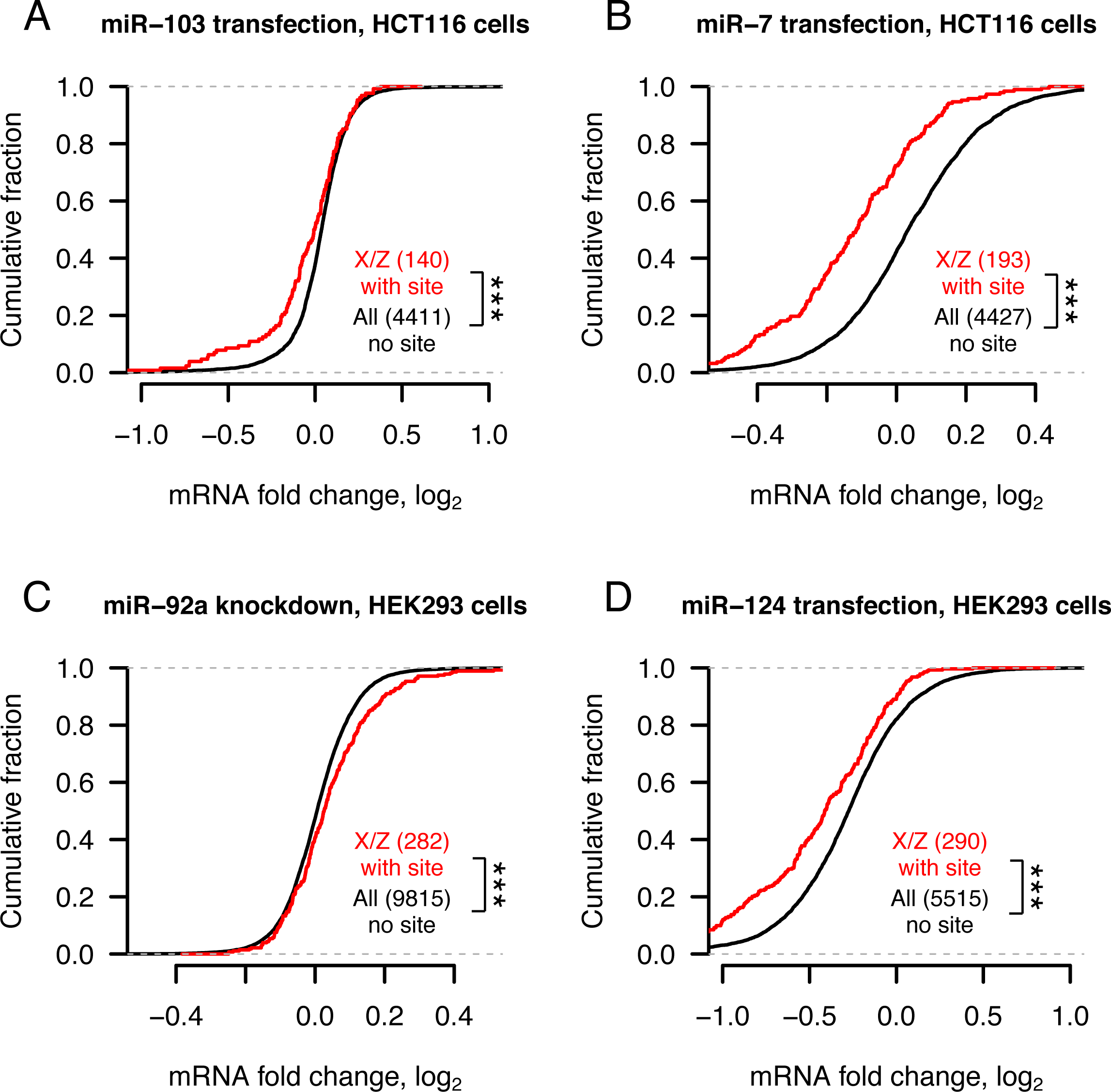
Gene expression changes following transfection or knockdown of additional miRNAs in human HCT116 or HEK293 cells. *** p < 0.001, two-sided Kolmogorov-Smirnov test.

**Supplemental Figure S17:**
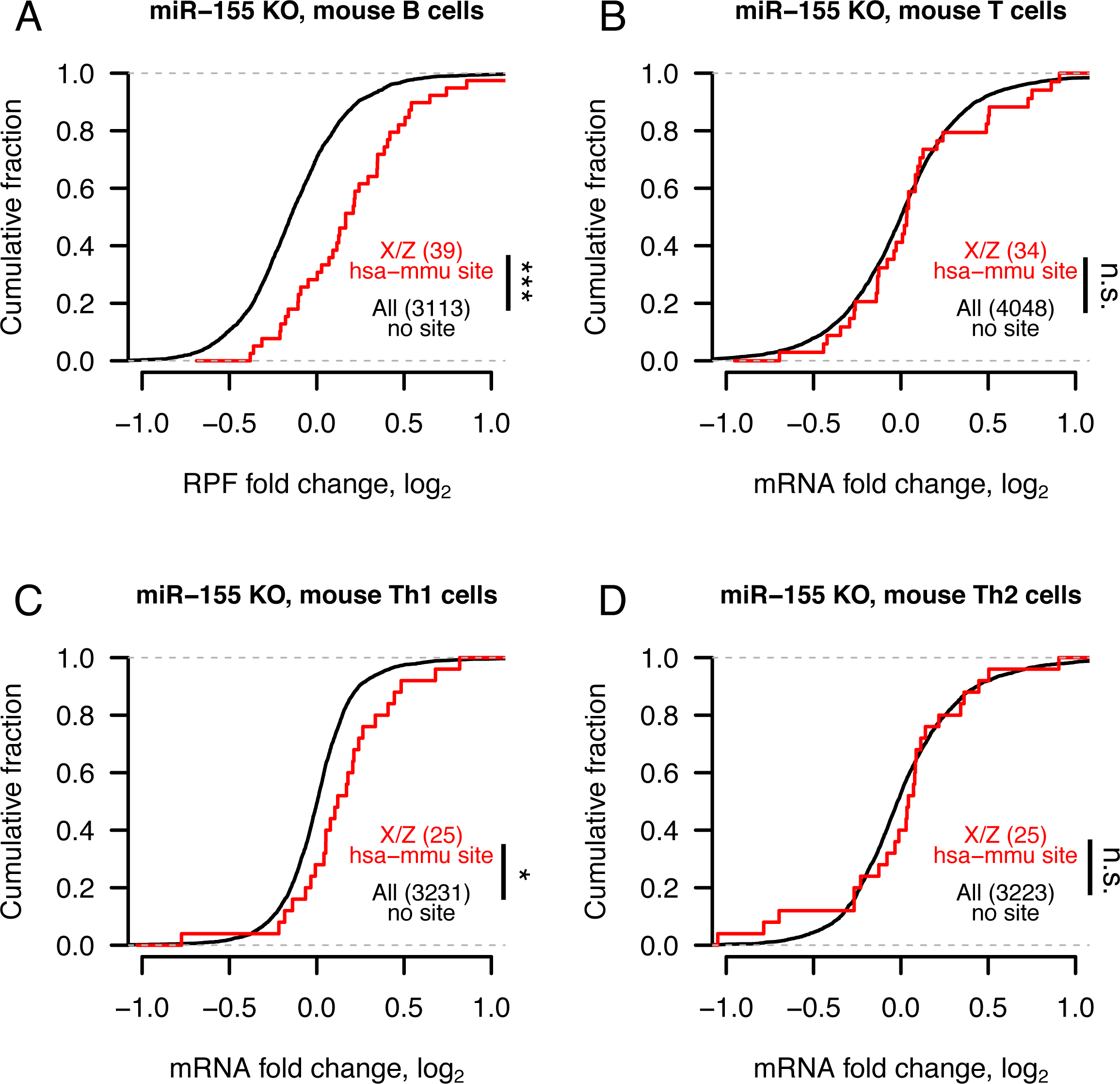
Changes in mRNA stability and translational efficiency and gene expression following miR-155 knockout in mouse immune cells. In each case, mouse orthologs of X- or Z-linked genes containing a human-mouse-conserved (hsa-mmu) miR-155 site were compared to mouse genes containing only nonconserved miR-155 sites. * p < 0.05, *** p < 0.001, two-sided Kolmogorov-Smirnov test.

**Supplemental Figure S18:**
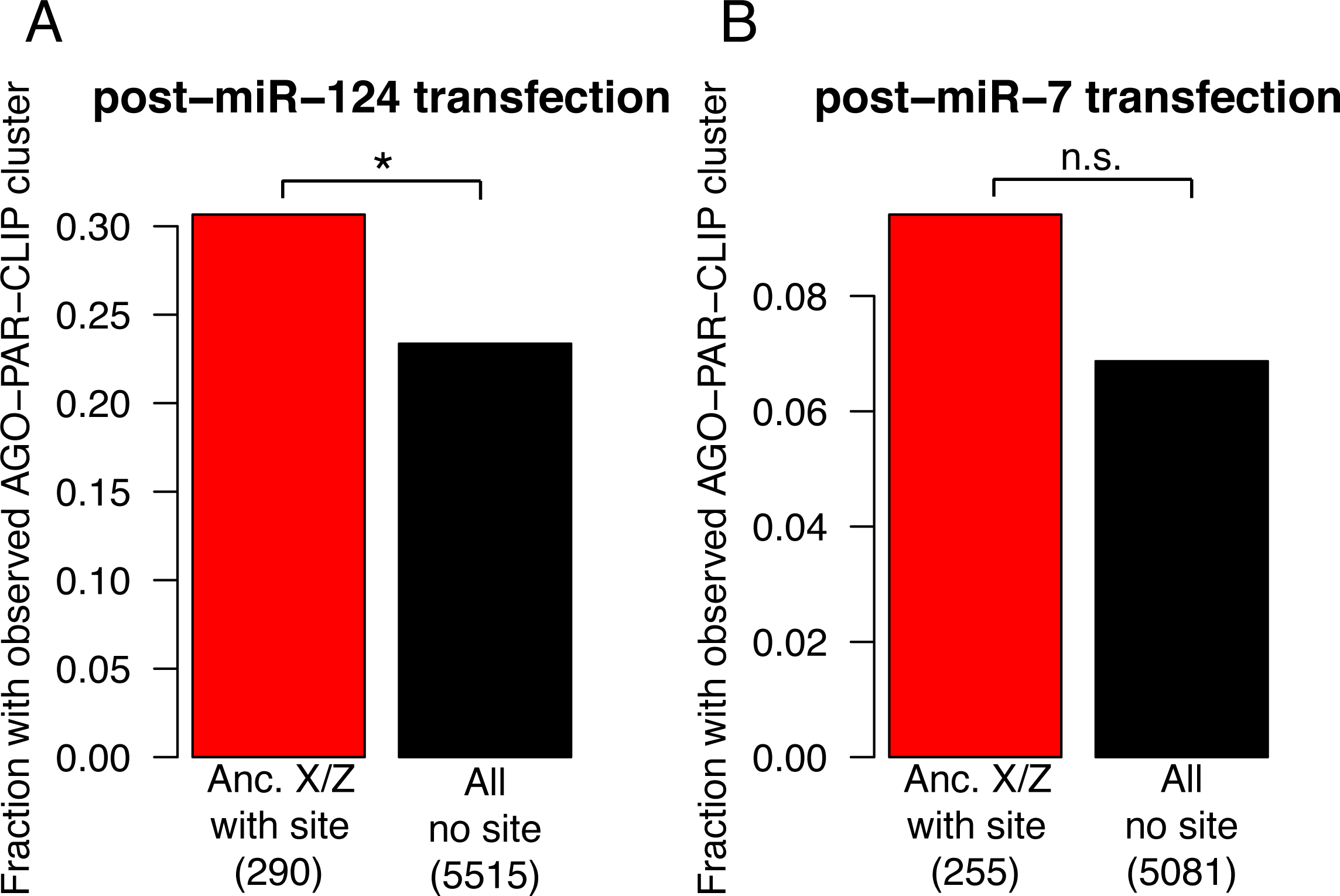
Argonaute binding measured by high-throughput crosslinking-immunoprecipitation (CLIP) following miRNA transfection in HEK293 cells. * p < 0.05, two-sided Fisher’s exact test.

